# Path2Omics: Enhanced transcriptomic and methylation prediction accuracy from tumor histopathology

**DOI:** 10.1101/2025.02.26.640189

**Authors:** Danh-Tai Hoang, Eldad D. Shulman, Saugato Rahman Dhruba, Nishanth Ulhas Nair, Ranjan K. Barman, H. Lalchungnunga, Omkar Singh, MacLean P. Nasrallah, Eric A. Stone, Kenneth Aldape, Eytan Ruppin

## Abstract

Precision oncology is becoming increasingly integral to clinical practice, demonstrating notable improvements in treatment outcomes. While molecular data provide comprehensive insights, obtaining such data remains costly and time-consuming. To address this challenge, we developed Path2Omics, a deep learning model that predicts gene expression and methylation from histopathology for 23 cancer types. Path2Omics was trained on 20,497 slides (9,456 formalin-fixed and paraffin-embedded (FFPE) and 11,041 fresh frozen (FF)) from 8,007 patients across 23 The Cancer Genome Atlas cohorts. When tested on FFPE slides, the most readily available format in clinical pathology practice, the integrated model outperformed its individual FF and FFPE components, robustly predicting nearly 5,000 genes on average, approximately five times more than our recently published DeepPT model. Externally evaluated on seven independent cohorts, Path2Omics robustly predicted the expression of approximately 4,400 genes, yielding a 30% increase over the FFPE model alone. Finally, we demonstrate that the inferred gene expression is nearly as effective as the actual values in predicting patient survival and treatment response. These results lay the basis for using Path2Omics to advance precision oncology from histopathology slides in a speedy and cost-effective manner.

**Statement of significance:** Path2Omics is a deep learning model that accurately predicts gene expression and methylation from histopathology slides across 23 cancer types. Unlike existing approaches that rely solely on FFPE slides for training, Path2Omics leverages both FFPE and FF slides by constructing two separate models and integrating them. Downstream analyses show that the inferred values from Path2Omics are nearly as effective as actual values in predicting patient survival and treatment response.

## Introduction

Pathology images have been routinely used in clinical practice since the early 1900s, ranging from tumor detection to subtype classification and prognosis (van den Tweel and Taylor 2010). Recent advances in artificial intelligence models, combined with the increasing availability of digital pathology, have enabled the ability of performing these tasks in an automatic manner, even without pathologist annotations (Teichmann et al. 2022; Courtiol et al. 2019). Furthermore, deep learning frameworks have demonstrated potential in predicting molecular signatures, including genetic mutations (Coudray et al. 2018; Kather et al. 2019; Fu et al. 2020; Qu et al. 2021; Schaumberg, Rubin, and Fuchs 2018; Tsou and Wu 2019; Chang et al. 2018; Kim et al. 2019; M. Chen et al. 2020; Ghaffari Laleh et al. 2022; Nasrallah et al. 2023), bulk mRNAseq expression (Y. Wang et al. 2021; Schmauch et al. 2020; Alsaafin et al. 2023; Hoang, Dinstag, et al. 2024), bulk DNA methylation (Hoang, Shulman, et al. 2024) and spatial mRNAseq expression (B. He et al. 2020; Monjo et al. 2022; Levy-Jurgenson et al. 2020; Shulman et al. 2024; C. Wang et al. 2025). Additionally, deep learning has been applied to predict patient survival (Mobadersany et al. 2018; Cheng et al. 2017; Beck et al. 2011; Bulten et al. 2020; Courtiol et al. 2019; Boehm et al. 2022; Y. Chen et al. 2023), and treatment response from histopathology (Xiangxue Wang et al. 2022; Hu et al. 2021; Johannet et al. 2021; F. Zhang et al. 2020).

With respect to the types of image preparation, previous studies have predominantly utilized Formalin-Fixed Paraffin-Embedded (FFPE) slides, also known as *diagnostic slides*, as they are the standard in clinical practice, thanks to their high quality and their central place in the diagnostic workflow of pathologists. To the best of our knowledge, no study has yet leveraged Fresh Frozen (FF) slides, also termed *tissue slides*, to develop deep learning models for predicting gene expression and methylation from slides. Our motivation to explore the predictive potential of FF slides stems from the observation that, in TCGA (a major resource for training predictive omics models from pathology slides), the available FF slides indeed correspond to the adjacency sections (“top-section” and “bottom-section”) of the slide used for RNA and DNA sequencing (“middle-section”). In difference, FFPE slides, although derived from the same tumor specimen, are not always in close proximity to the actual tissue used for gene expression and methylation analyses (Cooper et al. 2018). Consequently, there is a tradeoff between image quality and matching the molecular measurements when using only one type of slide from TCGA to build models for predicting gene expression and methylation. In this study, we aim to assess the predictive performance of each slide type and, more importantly, investigate whether integrating FF and FFPE slides enhances gene expression and methylation prediction. Furthermore, we aim to evaluate if the predicted values could be used to effectively predict patient survival and response to cancer therapies.

To address this challenge, we developed Path2Omics, a deep learning framework that integrates FFPE and FF slides to predict gene expression and methylation for 23 cancer types. Specifically, for each prediction task and cancer type, we trained two separate modes: one based on the FFPE slides, called “*FFPE model*”, and another based on FF slides, called “*FF model*”. Our final *“integrated model”* combines both predictions from the FFPE model and FF model. To study the potential application of the predicted gene expression and methylation, we subsequently developed models to predict patient survival and treatment response from their inferred values. Our results demonstrate that the model performance derived from their slide inferred values is remarkably comparable and similar to that obtained from the actual measured omics values.

## Results

### Study overview

We employed a deep learning framework that combines both FFPE and FF slides to accurately predict gene expression and methylation and subsequently predict patient survival and treatment response from the predicted gene expressions. Specifically, we analyzed 20,497 slides (9,456 FFPE and 11,041 FF) and their matched gene expression and methylation profiles from 8,007 patients across 23 TCGA cancer types to develop our models. For each cancer type, we trained two separate models, an *FFPE model* based on FFPE slides and an *FF model* based on FF slides. The number of slides and patients for each cancer type is provided in **Extended Data Table 1** (for gene expression) and **Extended Data Table 2** (for methylation). These models, and the model that integrates both, termed the *integrated model*, were first evaluated through cross-validation on the 23 TCGA cancer types. They were then further tested on seven external datasets, comprising 1,323 slides (1,163 FFPE and 160 FF) (**Extended Data Table 3**). As we shall show, the integrated model, which computes the mean scores of the FF and FFPE models, outperforms each individual model. To further evaluate model performance, we developed models to predict patient survival from the predicted gene expression for 12 TCGA cohorts. Finally, we built models to predict patient response to chemotherapy and immunotherapy based on the inferred gene expression (see **Methods** for more details).

### Predicting gene expression from H&E slides

After training the models, their performance was evaluated by measuring the Pearson correlation between the actual and predicted expression values for each gene across all samples within each cohort. Approximately 18,000 genes were analyzed independently for each cancer type. We first assessed the performance of our models using a fivefold cross-validation strategy on each individual cancer type on the TCGA cohort, where the model was iteratively trained on a subset of 80% of patients and used to make predictions on the remaining 20%. We found that almost every gene had a positive correlation between the actual and predicted expression values. Specifically, the FFPE models achieved an average of median correlation of 0.27 across the 23 cancer types (**Extended Data Fig. 1,** gray). Interestingly, the FF models significantly outperformed the FFPE models, achieving an average of median correlation of 0.36 (**Extended Data Fig. 1,** pink). In measuring the number of *well-predicted* genes, defined as those with a correlation between the inferred and measured expression values across all patient samples above 0.4, the FFPE model achieved an average of 3,503 highly predicted genes across the 23 cancer types, ranging from 584 genes (UCEC) to 9,756 genes (TGCT) (**Fig. 2a**, gray). Remarkably, the performance of the FF models was approximately double that of the FFPE models, achieving an average of 7,299 well-predicted genes, ranging from 2,872 genes (GBM) to 13,344 genes (TGCT) (**Fig. 2a**, pink).

**Fig. 1.**
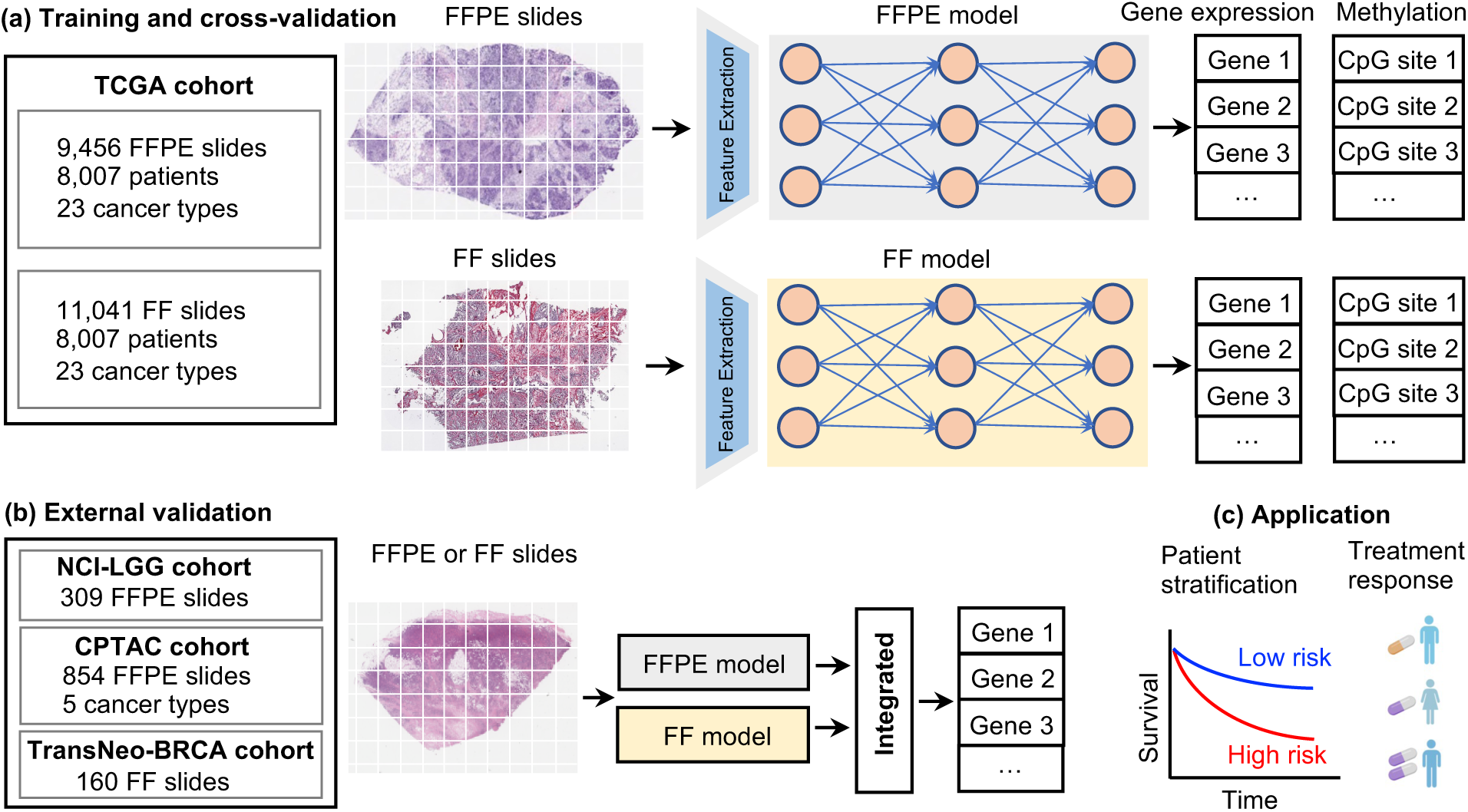
Overview of the computational workflow. **(a)** Path2Omics consists of two components: an FFPE model and an FF model. Both models share the same architecture and comprise three main units: image pre-processing, feature extraction, and regression. In the image pre-processing step, whole slide images were divided into tiles, followed by color normalization. In the feature extraction unit, the CTransPath digital pathology foundational model was used to encode each tile image into a 768-dimensional feature vector. Finally, in the regression unit, a multi-layer perceptron (MLP) was employed to process the extracted features and predict gene expression or methylation values. Each component was trained and cross-validated using the TCGA dataset, independently for each cancer type. A total of 20,497 slides (9,456 FFPE and 11,041 FF) and their matched gene expression and methylation profiles from 8,007 patients across 23 TCGA cancer types were used for training. **(b)** For external validation, we analyzed 1,323 slides (1,163 FFPE and 160 FF) from seven datasets obtained from three independent resources: in-house (NCI), CPTAC, and TransNeo. **(c)** To assess the clinical application of the inferred gene expression, we built models to predict patient survival and treatment response based on the predicted gene expressions.

**Fig. 2.**
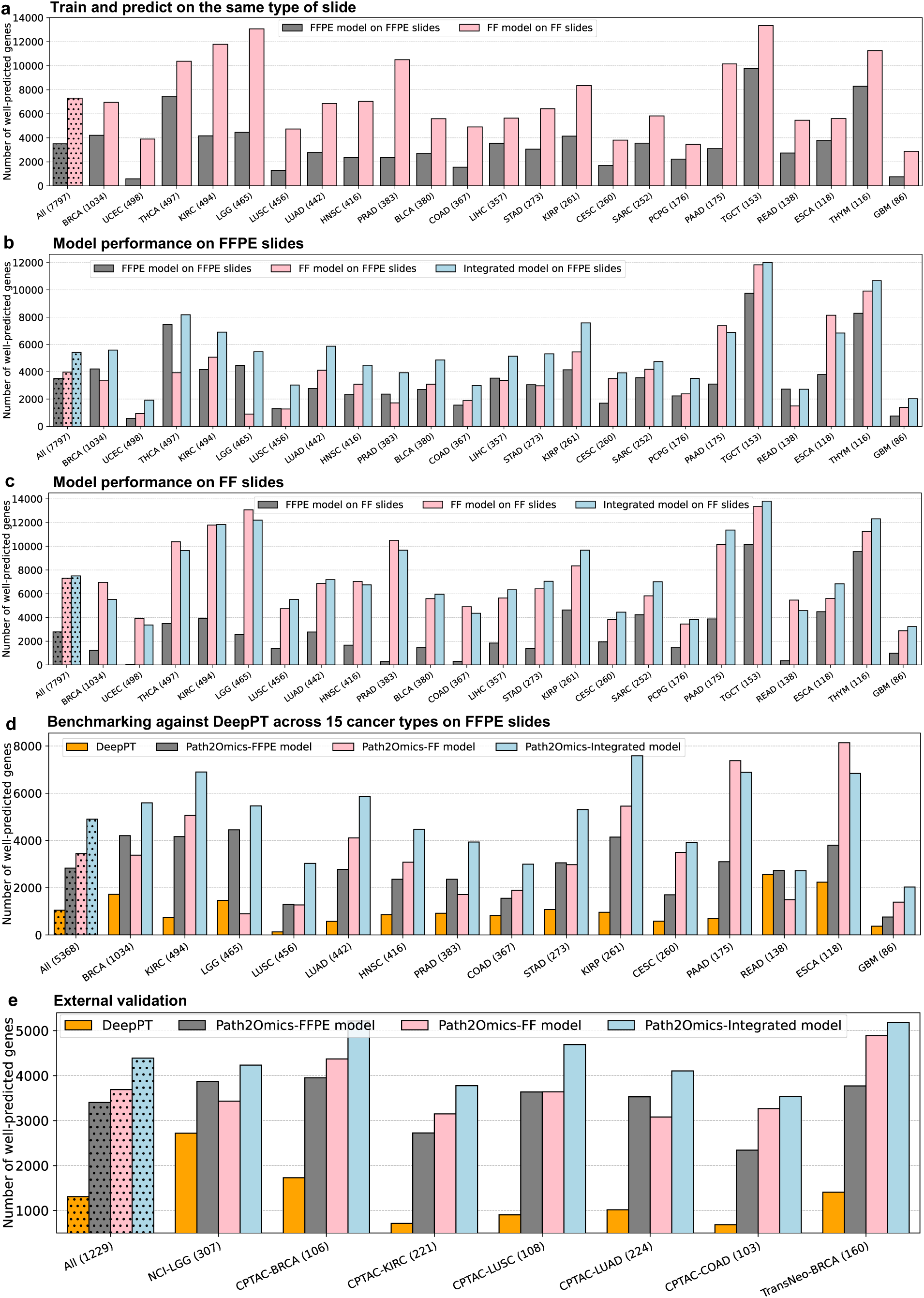
Path2Omics performance in predicting gene expression. **(a)** The number of well-predicted genes, defined as those with a Pearson correlation above 0.4 between predicted and actual expression values across the cohort samples, achieved by the Path2Omics-FFPE model on FFPE slides (gray), and the Path2Omics-FF model on FF slides (pink). The first bars represent the average across the 23 cohorts. The number of patients in each cohort is shown in parentheses. **(b)** The number of well-predicted genes achieved by the FFPE model (gray), FF model (pink) and Integrated model (light blue) when tested on FFPE slides across 23 TCGA cancer cohorts. The first bars represent the average across the 23 cohorts. **(c)** Similar plot to (**b**), but when tested on FF slides. **(d)** Benchmarking against the state-of-the-art approach, DeepPT. The number of well-predicted genes achieved by the DeepPT (orange), Path2Omics-FFPE model (gray), Path2Omics-FF model (pink) and Path2Omics-Integrated model (light blue) on FFPE slides from the 15 cancer types analyzed by both DeepPT and Path2Omics. **(e)** Similar plot to (**d**), but when tested on 7 external datasets, including FFPE and FF slides.

To evaluate the generalizability of each model to different types of slide preparations, we applied the pre-trained FF model to predict gene expressions on FFPE slides in the TCGA cohort and vice versa. Remarkably, the FF model generalized well to predicting the expression from FFPE slides, achieving an average of 3,973 well-predicted genes across the 23 cancer types (**Fig. 2b**, pink). Conversely, by applying the pre-trained FFPE model to predict gene expression on FF slides in the TCGA cohort, we obtained an average of 2,786 well-predicted genes (**Fig. 2c**, gray), approximately 30% lower than that achieved by applying the pre-trained FF model on FFPE slides (2,786 well-predicted genes versus 3,973 well-predicted genes). These results suggest that the pre-trained FF models generalized better than the pre-trained FFPE models on different types of slide preparations (**Fig. 2c**, light blue). Importantly, the Path2Omics integrated model, which combines predictions from the FFPE model and the FF model, outperformed each model individually, achieving an average of 5,417 well-predicted genes on the FFPE slides (**Fig. 2b**, light blue), representing an approximately 50% increase compared with the FFPE alone (5,417 versus 3,503 well-predicted genes).

Benchmarking against the current start-of-the-art approach, DeepPT (Hoang, Dinstag, et al. 2024), across the 15 cancer types analyzed by both DeepPT and Path2Omics on FFPE slides, the Path2Omics-FFPE model and Path2Omics-FF model achieved 2,830 and 3,449 well-predicted genes, respectively, approximately 3 times higher than the average of 1,047 well-predicted genes achieved by DeepPT across the different cancer types. Remarkably, the Path2Omics-Integrated model achieved 4,903 well-predicted genes, approximately 5 times higher than DeepPT (**Fig. 2d**). Of note, to ensure an apple-to-apple comparison, benchmarking was conducted exclusively on traditional FFPE slides, as analyzed by DeepPT.

To further evaluate the generalizability of our model, we applied the pre-trained FFPE and pre-trained FF models to predict gene expression profiles on 7 external datasets, consisting of 1,323 slides, including 1,163 FFPE slides and 160 FF slides. Quite strikingly, even though 6 of 7 datasets are composed of FFPE slides solely, the FF models achieved an average of 3,691 well-predicted genes across the seven datasets, outperforming the FFPE models, which achieved an average of 3,404 well-predicted genes. The integrated model achieved an average of 4,391 well-predicted genes on these external validation patient datasets, representing an approximately 30% increase compared to the traditional FFPE model (**Fig. 2e**).

Examining the overlap of the well-predicted genes obtained by the FFPE model and the FF model in each external cohort, we found that an average of 79% (ranging from 70% to 85%) of the genes achieved by the FFPE model were also achieved by the FF model. This finding once again highlights the generalizability of the FF model to FFPE slides (six out of seven external datasets are FFPE slides) (**Extended Data Fig. 2a**). Considering the well-predicted genes obtained by both the FFPE model and the FF model with those obtained by the integrated model, we found that 100% of these genes were also achieved by the integrated model (**Extended Data Fig. 2b**). Furthermore, considering the well-predicted genes obtained by either the FFPE model or the FF model, we found that an average of 92% (ranging from 87% to 96%) of these genes were also achieved by the integrated model (**Extended Data Fig. 2c**). The pronounced overlaps highlight the robustness of the integrated model, indicating that the integrated model not only retains the consistent performance of each component but also combines the strengths of both.

To identify cancer hallmarks (Hanahan and Weinberg 2011; Iorio et al. 2018) associated with well-predicted genes achieved by Path2Omics-integrated models, we performed pathway enrichment analysis across 23 TCGA cancer types. We observed similar patterns between FFPE and FF slides (**Fig. 3**). Specifically, four hallmarks, “tumor promoting inflammation”, “sustaining proliferative signaling”, “activating invasion and metastasis”, and “inducing angiogenesis” were consistently enriched across all 23 cancer types, whereas the remaining hallmarks were rarely enriched. The enrichment patterns for each cancer type are detailed in **Extended Data Fig. 3** (for FFPE slides) and **Extended Data Fig. 4** (for FF slides).

**Fig. 3.**
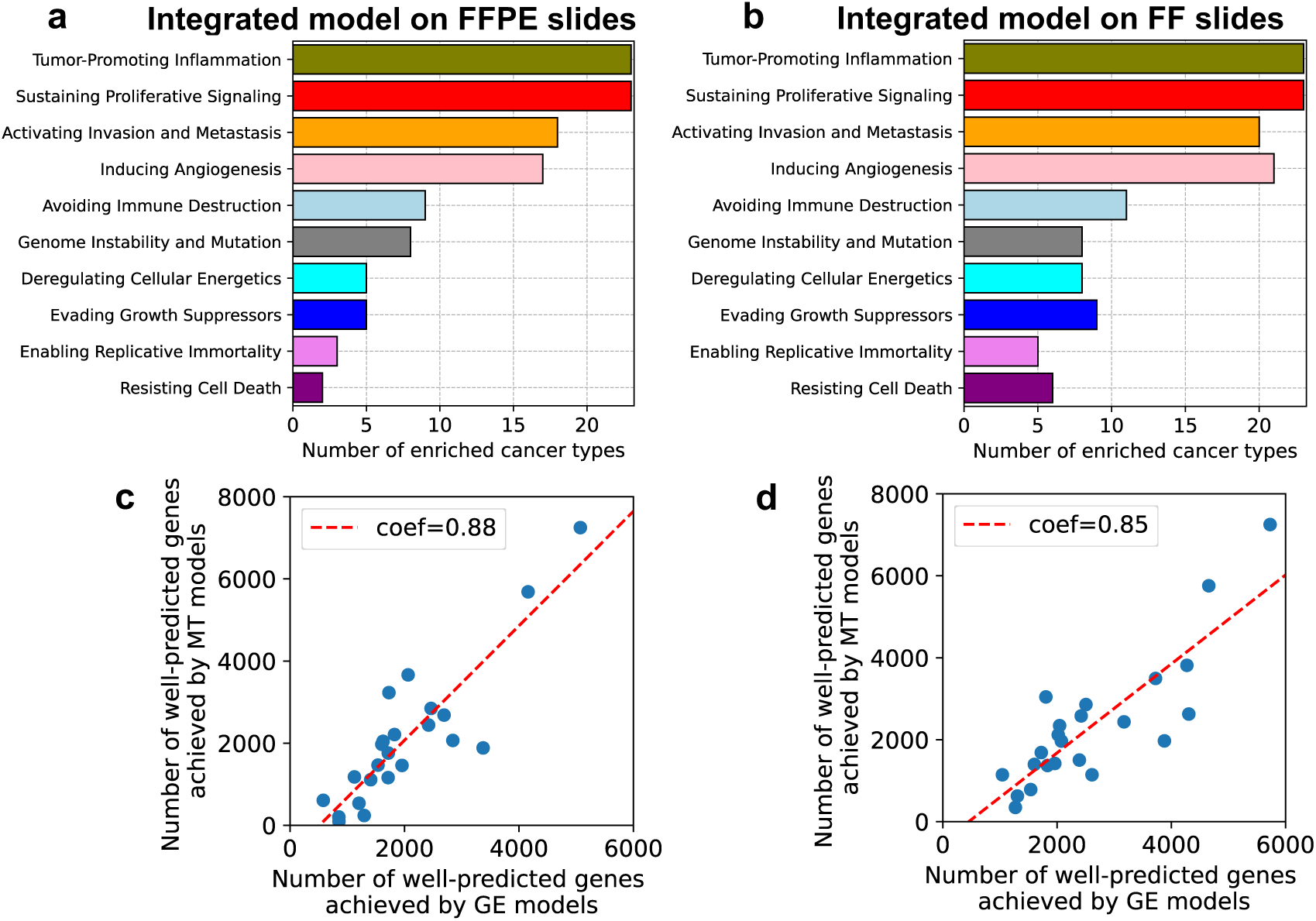
Cancer pathway enrichment analysis and the correlation between model performance in predicting gene expression and methylation. The number of cancer types in which each cancer hallmark was enriched with well-predicted genes, as achieved by running the Path2Omics-integrated models on either FFPE slides (**a**) or FF slides (**b**). The association between the number of well-predicted genes achieved by Path2Omics integrated model in predicting gene expression models (GE models, x-axis) and in predicting methylation (MT models, y-axis) across 23 TCGA cancer types, when tested on FFPE slides (**c**) and FF slides (**d**). Each data point represents one cancer type.

### Predicting DNA methylation from H&E slides

Using the same computational pipeline applied to gene expression above, we developed models to predict DNA methylation beta values from tumor H&E slides. For each cancer type, we selected the 40,000 most highly variable CpG probes for analysis. Consistently, our results showed that the FF models outperformed and generalized better than FFPE models across different types of slide preparations. Furthermore, the integrated models demonstrated improved performance over each individual component.

Specifically, when tested on the same type of slide preparation, the FFPE models achieved an average median correlation of 0.27, and an average of 9,223 well-predicted CpG probes (as before, correlations > 0.4) across the 23 cancer types (**Extended Data Fig. 5** and **Extended Data Fig. 6a,** gray). In comparison, the FF models achieved an average median correlation of 0.33 and an average of 13,194 well-predicted CpG probes (**Extended Data Fig. 5 and Extended Data Fig. 6a,** pink). When tested on different types of slide preparations, the FF models observed an average of 8,669 well-predicted probes on FFPE sides (**Extended Data Fig. 6b,** pink), outperforming the FFPE model, which observed an average of 7,759 well-predicted probes on FF slides (**Extended Data Fig. 6c**, gray). Once again, the integrated model outperformed the individual models, achieving an average of 12,272 well-predicted probes on FFPE slides (**Extended Data Fig. 6b**, light blue) and 14,544 well-predicted probes on FF slides (**Extended Data Fig. 3c**, light blue). This corresponds to a 33% increase (12,272 versus 9,223) and 87% (14,544 versus 7759) compared to the traditional FFPE models, respectively.

To compare the performance of Path2Omics-integrated models in predicting gene expression and methylation, we aggregated both measured and predicted methylation beta values from the CpG site level to the gene level, focusing on common genes included in both tasks. Across 23 cancer types, we observed a strong association between the number of well-predicted genes achieved by the two models, with a Pearson correlation of 0.88 for FFPE slides (**Fig. 3c**) and 0.85 for FF slides (**Fig. 3d**). This finding suggests that in cohorts where Path2Omics performs well in predicting gene expression, it also performs well in predicting methylation.

### Predicting patient survival from the predicted gene expression

To evaluate the potential translational value of the predicted gene expression from Path2Omics, we developed models to predict patient overall survival using the latter. Specifically, we focused on twelve TCGA cancer type cohorts with sufficient sample sizes (over 100 samples) (**Extended Data Table 3**). For each cancer type, we employed multivariate Cox-regression models incorporating the inferred gene expression of robustly predicted genes along with patient demographic information (sex and age) to predict patient overall survival (**Methods**). The models were trained and tested using cross-validation.

The models successfully stratified patients into high-risk and low-risk groups in 11 out of 12 cohorts (p-value < 0.05), except for LUSC (p-value = 0.112) (**Fig. 4a**, “Predicted GE”, left panels). For comparison, we independently developed analogous survival prediction models based on the actual gene expression and patient demographics. We found that the two models, one based on the inferred gene expression (**Fig. 4a**, “Predicted GE”, left panels), and the other based on the actual gene expression (**Fig. 4a**, “Actual GE”, right panels), produced nearly identical Kaplan-Meier curves across all the 12 cohorts.

**Fig. 4.**
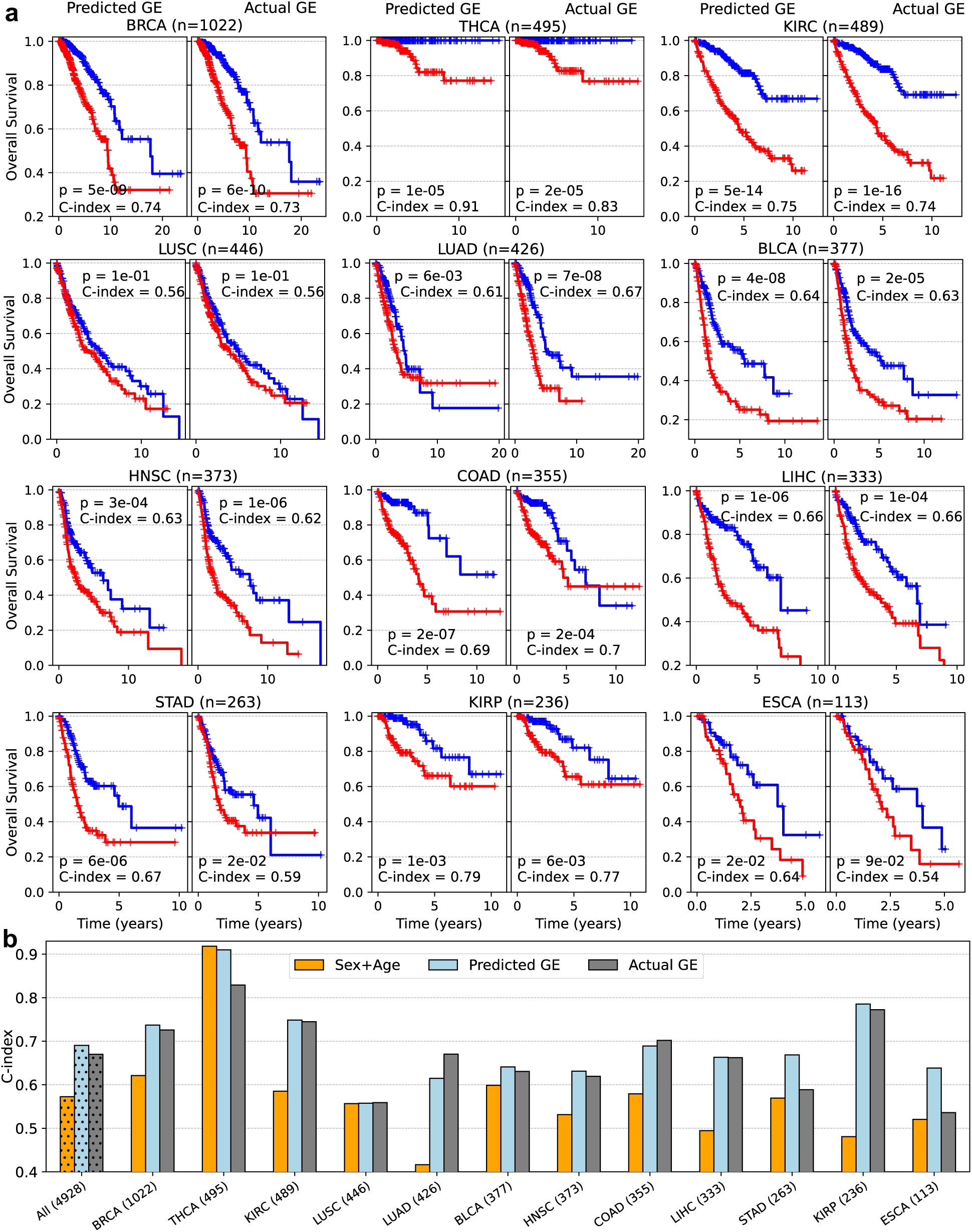
Model performance in predicting patient survival based on inferred and measured gene expression. **(a)** Kaplan-Meier curves generated by model using predicted gene expression and patient demographics (sex, age) (“Predicted GE”, left panels) compared with those from the model using actual gene expression and patient demographics (“Actual GE”, right panels) across 12 cancer cohorts. In each cohort, high-risk (red) and low-risk (blue) groups were stratified based on the median risk score generated by each model. P-values were calculated using a two-sided log-rank test. **(b)** Concordance index achieved by the “Predicted GE” model (light blue) compared with the “Actual GE” model (gray) and a control model using only patient sex and age (orange). The first bars represent the average C-index across the 12 cohorts. The number of patients in each cohort is indicated in parentheses.

To quantitatively assess model performance, we computed the concordance index (C-index), which measures how well the predicted risk scores align with observed outcomes for pairs of patients. The “Predicted GE” model achieved a mean high C-index of 0.69 across the 12 cohorts (**Fig. 4b**, light blue bars). This performance was similar to that of the “Actual GE” model (average C-index of 0.67, **Fig. 4b**, gray bars), and remarkably outperformed the model using only the patient sex and age (average C-index of 0.57, **Fig. 4b**, orange bars).

Furthermore, analyzing the overlap between the high-risk patient group identified by the “Predicted GE” and “Actual GE” models in each cohort, we found that an average of 71% of patients across the 12 cohorts (ranging from 63% in HNSC to 82% in KIRC) were consistently classified as high risk by both models (**Extended Data Fig. 7**). The strong concordance of the “Predicted GE” model compared with the “Actual GE” model further validates the potential clinical relevance of the predicted gene expression by Path2Omics.

Similarly, we built survival prediction models using inferred methylation data and compared them with models based on actual methylation data. Once again, we observed that the Kaplan-Meier curves derived from models using inferred methylation were very similar to those obtained using actual methylation across all the 12 cohorts (**Extended Data Fig. 8**). Furthermore, the high-risk patient groups identified by the two models overlapped substantially, with an average of 69% of patients across the cohorts (ranging from 62% in LUSC to 80% in KIRC) (**Extended Data Fig. 9**).

### Predicting breast cancer patient response to chemotherapy and trastuzumab treatments from Path2Omics inferred gene expression

To further validate the potential clinical value of Path2Omics inferred gene expression, we reconstructed models for predicting patient response to cancer therapies using those values. Specifically, we utilized four distinct classical machine-learning algorithms to develop an ensemble model that combines their prediction scores (**Methods**). Our models were trained and tested through cross-validation on two breast cancer cohorts: one with HER2-negative patients treated with chemotherapy (93 patients, referred to as the *chemotherapy cohort*) and another with HER2-positive patients treated with chemotherapy plus trastuzumab (61 patients, referred to as the *trastuzumab cohort*) (Sammut et al. 2022).

In the chemotherapy cohort, the *Predicted GE* model based on predicted gene expression achieved an AUC of 0.82, an accuracy of 0.76 and an odds ratio of 7.6 (**Fig. 5a-c**, left column, dark blue) in predicting patient response. This performance was lower than that of the “*Actual GE*” model based on actual gene expression, which achieved an AUC of 0.87, an accuracy of 0.78, and an odds ratio of 8.1 (**Fig. 5a-c**, left column, gray). However, both models markedly outperformed the *Direct* model, which predicts treatment response directly from slides, without using the inferred gene expression data as an intermediary. The *Direct* model achieved an AUC of 0.74, an accuracy of 0.67, and an odds ratio of 5.7 (**Fig. 5a-c**, left column, orange).

**Fig. 5.**
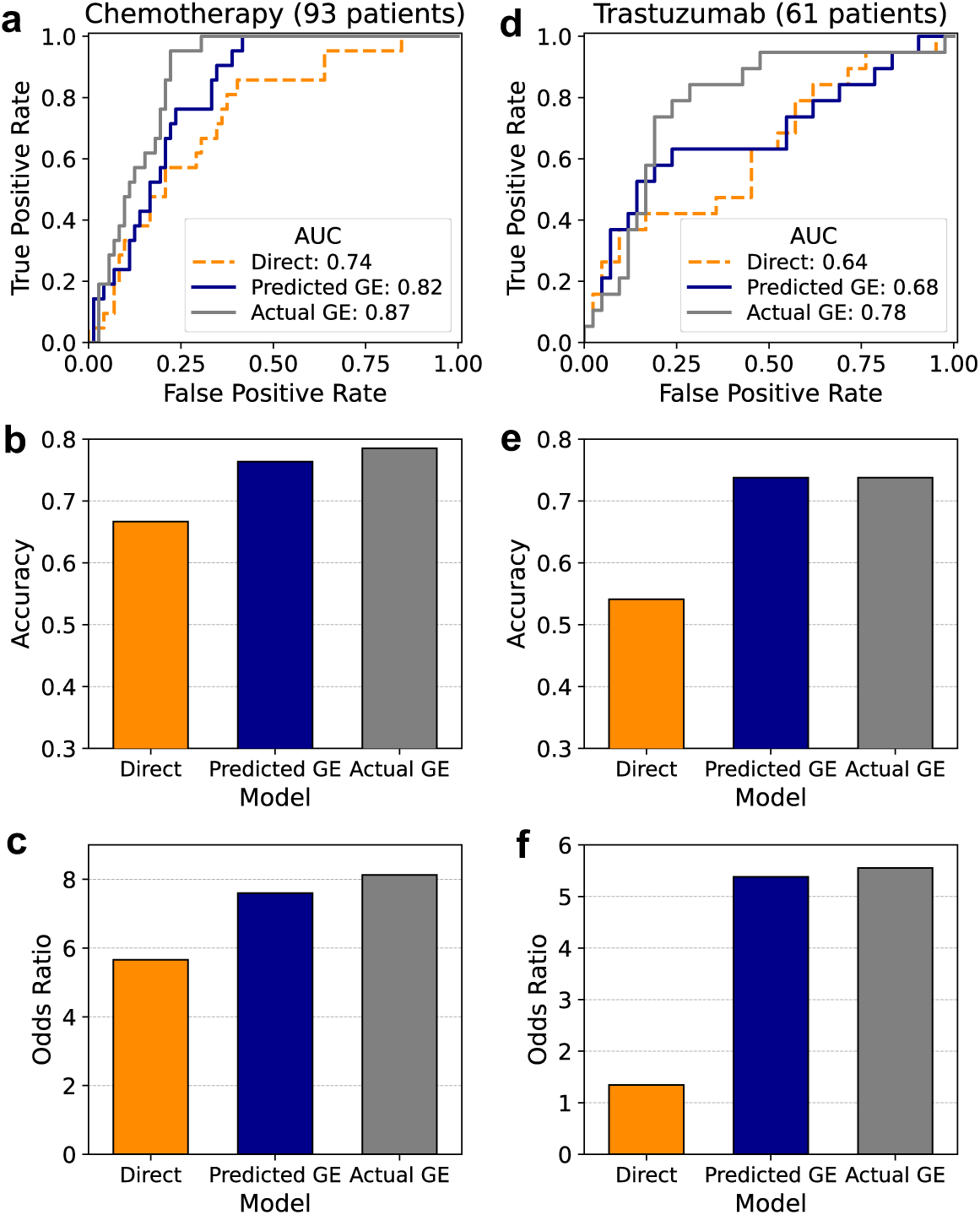
Model performance in predicting patient response to cancer therapy. AUC (**a**), accuracy (**b**), and odds ratio (**c**) for the chemotherapy cohort (left column) and trastuzumab cohort (right column) are shown for three models: the “Direct” model that predicts response directly from pathology images (orange) without intermediate step of predicted gene expression, the “Predicted GE” model that predicts response from predicted gene expression (dark blue), and the “Actual GE” model that predicts response from actual gene expression (gray).

In the “trastuzumab” cohort, the *Predicted GE* model based on the inferred gene expression achieved an AUC of 0.68, an accuracy of 0.74, and an odds ratio of 5.4 (**Fig. 5a-c**, right column, dark blue). Once again, these results were somewhat less than those obtained by the model based on actual gene expression (*Actual GE* model), which achieved an AUC of 0.78, an accuracy of 0.74, and an odds ratio of 5.6 (**Fig. 5a-c**, right column, gray). However, consistently, both models quite markedly outperformed the *Direct* model, which achieved an AUC of 0.64, an accuracy of 0.54, and an odds ratio of 1.35 (**Fig. 5a-c**, right column, orange). These results underscore Path2Omics ability to predict patient response to cancer therapies directly from the pathology slides, presenting a viable (even if somewhat less accurate), fast and low-cost alternative to precision oncology omics-based predictors.

## Discussion

We present Path2Omics, an integrated deep learning model that predicts gene expression and methylation from pathology slides. We demonstrate that Path2Omics markedly improves upon existing methods for predicting gene expression and methylation from whole slide images, including our own previously described algorithm (Hoang, Dinstag, et al. 2024). Additionally, we show that the predicted gene expression and methylation data are as effective as their actual data for predicting cancer patient survival across many TCGA datasets. In the prediction of breast cancer patient treatment response to chemotherapy and trastuzumab, we find that predictors based on the measured gene expression performs best, but predictors based on the inferred expression approach these accuracy levels, and we find them to outperform ‘direct’ predictors based on the pathology slide images as is (i.e., without inferring the gene expression data).

Over the last few years, several approaches have been developed to predict gene expression from pathology slides. As demonstrated in (Hoang, Dinstag, et al. 2024), our recently published DeepPT outperformed previous approaches in predicting bulk gene expression, including HE2NRA (Schmauch et al. 2020), SEQUOIA (Zheng et al. 2023), and tRNAsformer (Alsaafin et al. 2023). Furthermore, a recent independent benchmarking study recently showed that DeepPT outperformed other approaches in predicting spatial gene expression as well (C. Wang et al. 2025), comparing its performance to iStar (D. Zhang et al. 2024), Hist2ST (Zeng et al. 2022), DeepSpaCE (Monjo et al. 2022), HisToGene (Pang, Su, and Li 2021) and ST-Net (B. He et al. 2020). Two key differences that distinguish DeepPT from other existing approaches were its inclusion of an auto-encoder layer to compress 2,048 ResNet50 features into a lower-dimensional 512-vector, to avoid overfitting and its application of ensemble learning by averaging predictions across 25 models. Path2Omics builds upon DeepPT but introduces key and important modifications. Instead of using ResNet50 pre-trained on natural images from the ImageNet dataset (K. He et al. 2016), Path2Omics incorporates CTransPath, a pathology foundation model pre-trained on pathology slide images. Additionally, the auto-encoder unit in DeepPT was removed because the CTransPath-derived vector has only 768 dimensions obtained exclusively from pathology slide images, eliminating the need for further dimensionality reduction. Recent studies have demonstrated that foundation models show remarkably improved performance for computational pathology tasks (Xiyue Wang et al. 2024, 2022; Xu et al. 2024; R. J. Chen et al. 2024). Consistently, the number of robustly predicted genes by Path2Omics-FFPE model is 3 times higher than DeepPT when both models were trained and tested on FFPE slides from 15 TCGA cancer types. Moreover, the key innovation underpinning the substantial performance improvement by Path2Omics compared to DeepPT is that, unlike existing approaches that rely solely on FFPE slides for training, Path2Omics combines both FFPE and FF slides to build two separate models and then integrate them. As a result, the latter model achieves nearly 5,000 well-predicted genes, five times better than DeepPT even when tested on FFPE slides across the 15 cancer types. Beyond high accuracy in the prediction of gene expression, Path2Omics extends our previous work, DEPLOY, which predicted methylation from slides in brain cancer only (Hoang, Shulman, et al. 2024), to encompass 23 TCGA cancer types. Consistently, we found that the integrated model improved performance over the FF and FFPE individual models and over DEPLOY.

FFPE slides and FF slides are two common methods used in tissue sample preparation for histological analysis and molecular studies. FFPE slides provide excellent preservation of tissue architecture, making them widely used for histological analysis and routine clinical diagnostics by pathologists. However, formalin fixation may alter nucleic acids and proteins, potentially complicating downstream molecular analyses. In contrast, FF slides preserve the native state of biomolecules like proteins and nucleic acids and are widely used for RNA and DNA sequencing. Surprisingly, the deep learning model based on FF slides outperforms and generalizes better than the one based on FFPE model, even when applied to a dataset composed only of FFPE slides. While the reasons for this observation may be complex, these findings may be at least partially attributable to the fact the sequencing data was derived from locations that are more aligned to the FF slides than to FFPE slides. Nevertheless, it is intriguing that deep learning models are still able to extract a wealth of molecular information from FF slides, which could be challenging for pathologists. Most importantly, the integration of FF slides into the model training remarkably improves the accuracy of transcriptomic and methylation predictions from tumor histopathology, even when the model is tested exclusively on FFPE slides. The frozen section procedure, which involves rapidly freezing and analyzing tissue, is already used intraoperatively to determine whether a sample is tumor or normal, guiding surgical decisions in real time. AI-driven analysis of fresh frozen slides could enhance this process by improving accuracy, reducing variability, and integrating molecular insights, ultimately refining surgical and therapeutic strategies in oncology.

Although this work shows promising results, it has some limitations that should be addressed in future research. First, the use of the pre-trained FF model to predict on FFPE slides and vice versa was conducted directly, without any modifications or fine-tuning. Future work could explore using domain adaptation models such as generative adversarial networks (GANs) (Jose et al. 2021; Falahkheirkhah et al. 2022; Ozyoruk et al. 2022) to reduce the gap between the two types of slides. Second, the downstream analysis focused primarily on predicted gene expression data. Further studies could explore the application of predicted methylation data, for instance, in cancer subtype classification, as demonstrated in our previous work (Hoang, Shulman, et al. 2024).

Overall, our results provide strong evidence that integrating two different types of slide preparations while training gene expression and methylation predictors could markedly enhance their predictive performance, even when only one slide type is available during the prediction phase. Importantly, we show that downstream tasks, such as survival prediction and response to therapy, show high accuracy in our model, which has potential applications for clinical practice. As one example, a limitation of omics technologies in the clinic is the substantial time and cost involved. A model such as ours provides rapid initial insights that could potentially be used to provide clinically important information in situations where omics testing may not be feasible or available for patient populations. We hope that these findings will accelerate efforts to incorporate both FFPE and FF slides in developing such frameworks, ultimately making precision oncology more accessible in a timely manner.

## Methods

### Data collection

The Cancer Genome Atlas (TCGA) data, including histopathology slide images paired with bulk gene expression and DNA methylation, was downloaded from the Genomic Data Commons Data Portal (https://portal.gdc.cancer.gov/). We focused on the 23 cancer types with sample sizes sufficient for model training. For consistency, only patients with both Formalin-Fixed Paraffin-Embedded (FFPE) and Fresh Frozen (FF) slides were included in model development and evaluation. This selection strategy resulted in a total of 20,497 slides (9,456 FFPE and 11,041 FF) from 8,007 patients. Notably, some slides had only either matched gene expression or methylation, resulting in discrepancies between the number of slides and the number of patients for the two tasks. Specifically, for gene expression prediction, 19,963 slides (9,195 FFPE and 10,768 FF) from 7,797 patients were included, whereas for methylation prediction, 16,934 slides (8,159 FFPE and 8,775 FF) from 6,888 patients were used.

The in-house NCI-LGG dataset comprises 309 FFPE slides and their corresponding gene expression profiles from 307 patients, obtained from the Laboratory of Pathology at the National Cancer Institute (NCI).

The Clinical Proteomic Tumor Analysis Consortium (CPTAC) dataset includes FFPE slides paired with matched gene expression profiles. The pathology slides were downloaded from The Cancer Imaging Archive (TCIA; https://www.cancerimagingarchive.net), and the corresponding gene expression data were downloaded from the Genomic Data Commons Data Portal https://portal.gdc.cancer.gov/. For validation, five cancer types (BRCA, KIRC, LUSC, LUAD, and COAD) with sufficient samples from both the training cohort (TCGA) and the validation cohort (CPTAC) were selected. This selection resulted in 854 FFPE slides from 762 patients.

The TransNeo-Breast dataset consists of matched FF histopathology slides, RNAseq data, and treatment response information for 160 breast cancer patients. The slide images were generously shared by Dr. Sammut et al. (Sammut et al. 2022), and the RNAseq dataset was downloaded from the European Genome-Phenome Archive (EGA; accession number EGAS00001004582).

Clinical data for TCGA patient survival analysis, including overall survival, patient sex and age, were downloaded from (Liu et al. 2018).

Below, we present the Path2Omics computational framework for predicting gene expression. The same framework was applied to predict DNA methylation.

### Path2Omics computational framework

Path2Omics is an integrated model composed of two components: an FFPE model trained using FFPE slides and an FF model using FF slides. Both components share the same architecture and are composed of three main units.

<u>(1) Slide image processing:</u> To process the whole slide images (WSIs) for gene expression prediction models (or DNA methylation), Sobel edge detection (Kanopoulos, Vasanthavada, and Baker 1988) was applied to identify the tissue-containing areas within each slide. Due to the large size of the WSIs (ranging from 10,000 to 100,000 pixels in each dimension), they were divided at 20x magnification into non-overlapping tiles of 512 x 512 RGB pixels. Tiles where more than half of the pixels had a weighted gradient magnitude below a threshold (ranging from 10 to 20 depending on image quality) were removed. To minimize staining variation (heterogeneity and batch effects), Macenko’s method for color normalization was applied to the selected tiles (Macenko et al., 2023).
<u>(2) Feature extraction:</u> The pre-trained CTransPath model (Xiyue Wang et al. 2022) was used to extract features from the image tiles. During this process, each input tile is represented by a vector of 768 derived features, and each slide is represented as a matrix of size (n_tiles, 768), where n_tiles represents the number of selected tiles in the slide.
<u>(3) Multi-Layer Perceptron (MLP) regression:</u> To build a regression model that predicts gene expression (or DNA methylation) across whole genome using the 768 extracted features as input, an MLP module was used, consisting of three layers: (1) an input layer with 768 nodes, corresponding to the size of the extracted features; (2) a hidden layer with 512 nodes; and (3) an output layer with the number of nodes equal to the number of genes (or number of CpG sites) considered in the corresponding cancer cohort (approximately 18,000 genes, 40,000 CpG sites).

### Path2Omics training procedure

For each cancer type, we trained two separate models, one using FFPE slides with their matched gene expression (the “FFPE model”) and another using FF slides with their matched gene expression (the “FF model”). The training procedure of each model primarily followed the protocols outlined in DeepPT (Hoang, Dinstag, et al. 2024) and DEPLOY (Hoang, Shulman, et al. 2024). Briefly, we conducted a 5x5 nested cross-validation. In each outer loop, the entire set of samples for each cohort was split into a training set (80%) and a hold-out test set (20%). The training set was further subdivided into internal training and validation sets using another five-fold cross-validation, with the models trained and evaluated independently on each internal training-validation pair. Notably, the splits were performed at the patient level, ensuring that slides (both FFPE and FF) from the same patients were assigned to the same cross-validation set, thereby preventing information leakage between the training, validation and test sets.

To generate predictions for the TCGA dataset, we averaged the outputs from the five models, each trained on a different inner fold, for the samples in each outer fold. For the external validation sets (NCI, CPTAC, and TransNeo), the predictions from all 25 cross-validation models (five models for each of the five outer folds) were averaged to obtain the final prediction.

Each training round was stopped at a maximum of 500 epochs or earlier if the average correlation per gene between predicted and actual gene expression on the validation set did not improve for 50 consecutive epochs. The Adam optimizer was used with a mean squared error loss function, a learning rate of 0.0001, mini batches of 32 slides per step, and a dropout of 0.2 to prevent overfitting.

### Model for predicting patient survival

We developed and evaluated model performance using a five-fold cross-validation strategy. For each training fold, we first conducted univariate survival analysis to select the ten most predictive genes (or CpG sites). We then trained a multivariate Cox-regression model using these ten genes (or CpG sites) along with patient sex and age. The gene expression (or methylation) data and patient age were normalized to a range of 0-1 using scikit-learn’s “MinMaxScaler”, ensuring consistency across all features. Notably, feature selection and normalization were performed exclusively on the training samples to prevent data leakage, before applying to the test sets. For model evaluation, we used the C-index and Kaplan-Meier plots, both implemented in the Lifelines library.

### Model for predicting treatment response from gene expression

To predict patient response to cancer therapy based on predicted gene expression, we employed four independent classical machine-learning algorithms: logistic regression, support vector machine, *K*-nearest neighbor, and random forest. Within each training fold, we first normalized the gene expression values to a range of 0–1 using scikit-learn’s “MinMaxScaler”, ensuring consistency across all genes. Next, we selected the top 100 genes with the highest analysis of variance (ANOVA) *F-*values relative to response class using scikit-learn’s “SelectKBest” with the “f_classif” function. Each model was then trained using standard procedures. The final prediction score was calculated as the average of the prediction scores from the individual models. For model performance evaluation, we assessed the AUC using the implementation in Scikit-learn.

### Model for predicting treatment response directly from slides

The “Direct” model classifies patient response to cancer therapy directly from tumor slides, without the intermediate step of predicting gene expression. We used the same deep learning framework employed for predicting gene expression and methylation, except replacing the MLP regression unit with an MLP classification unit. Because the labels were available only at the slide level, the training followed a weakly-supervised learning approach in which all tiles from a given slide inherit the slide label (Coudray et al. 2018; Levy-Jurgenson et al. 2020; Fu et al. 2020; Kather et al. 2019; Ghaffari Laleh et al. 2022). After training, the slide-level prediction was computed by averaging the tile-level predictions for each slide.

### Gene expression processing

The gene expression data were originally provided as read counts. For our study, we selected only approximately 18,000 highly expressed genes for each cancer type independently. Within sample normalization was applied to minimize discrepancies in library size between experiments and batches (Hoang, Dinstag, et al. 2024).

### Ethics

Our study involves human data but does not involve direct interaction with human subjects. The two main cohorts, TCGA and CPTAC, are publicly available datasets.

### Randomizations

The training and test datasets in the cross-validation was split randomly.

### Statistical analysis

Statistical tests, including the method used and the sample sizes, are specified throughout the article.

### Implementation details

Analysis in this study was mainly performed in Python 3.9.7 with libraries including NumPy 1.20.3, Pandas 1.3.4, Scikit-learn 1.2.2 and MatPlotLib 3.4.3. Image processing including tile partitioning and color normalization was conducted with OpenSlide 1.1.2, OpenCV 4.5.4 and PIL 8.4.0. The feature extraction, MLP regression and MLP classification components were implemented using PyTorch 1.12.0. Classification machine-learning packages including logistic regression, support vector machine, *K*-nearest neighbor and random forest were deployed using Scikit-learn 1.2.2. Survival analysis was conducted using scikit-survival 0.23.1 and lifelines 0.27.0 packages.

### Code availability

The Path2Omics software tool will be available for academic research purposes upon acceptance of the publication at: https://doi.org/10.5281/zenodo.15016142

## Acknowledgements

This work was partially supported by the Intramural Research Program of the National Institutes of Health (NIH), National Cancer Institute (NCI), Center for Cancer Research (CCR).

## Author contributions

D.-T.H. developed the Path2Omics framework, discovered the use of fresh frozen slides for model development, and performed all analyses. E.D.S., S.R.D., N.U.N., R.B., M. P. N., E.A.S., and K.A. interpreted the results and provided feedback on the study. H.L, O.S., and K.A. provided the NCI-LGG external dataset. E.R. supervised the study. D.-T. H. and E.R. wrote the paper with assistance and feedback from all coauthors.

## Additional information

### Supplementary Information

**Supplementary Table 1.**
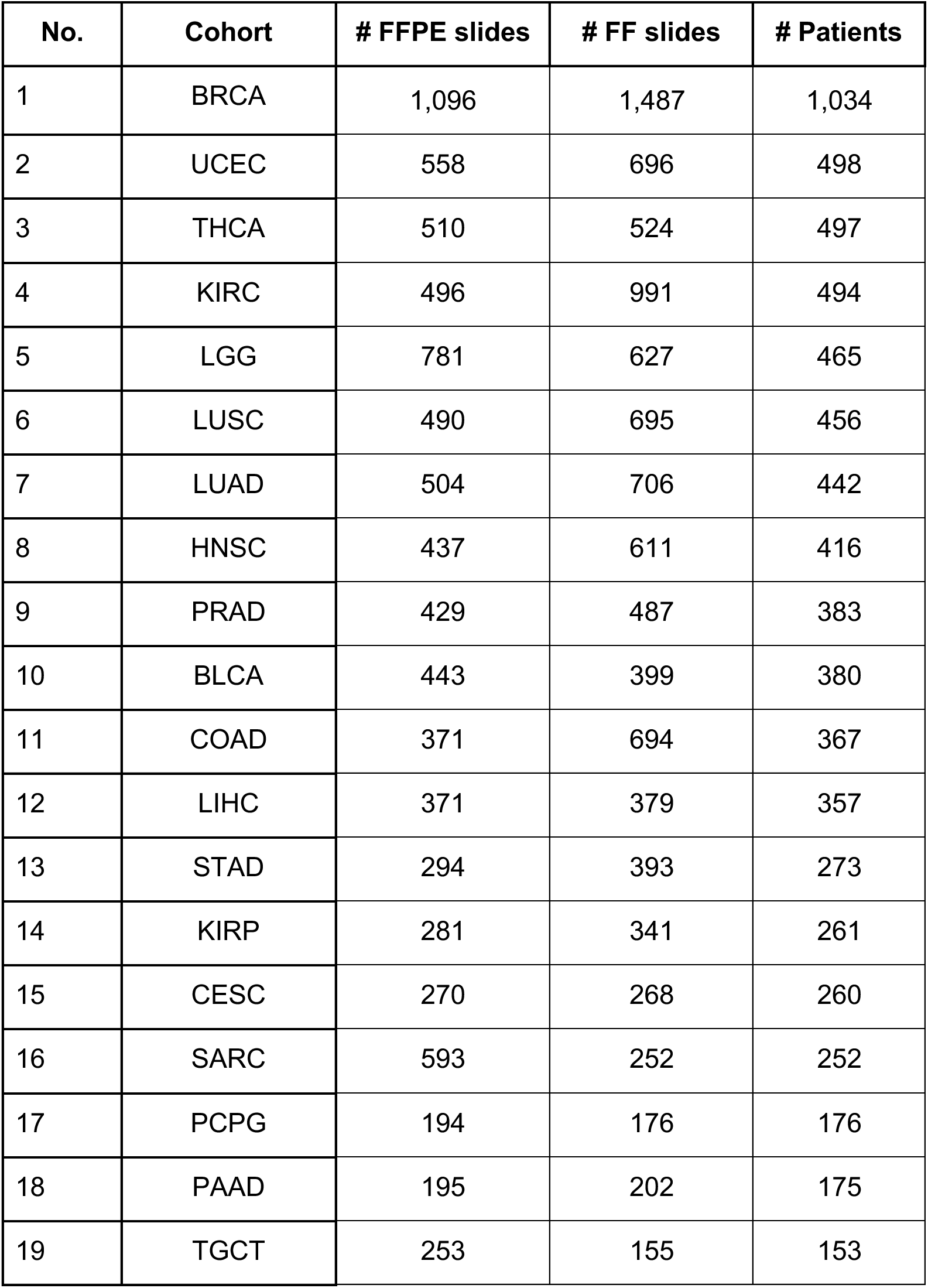
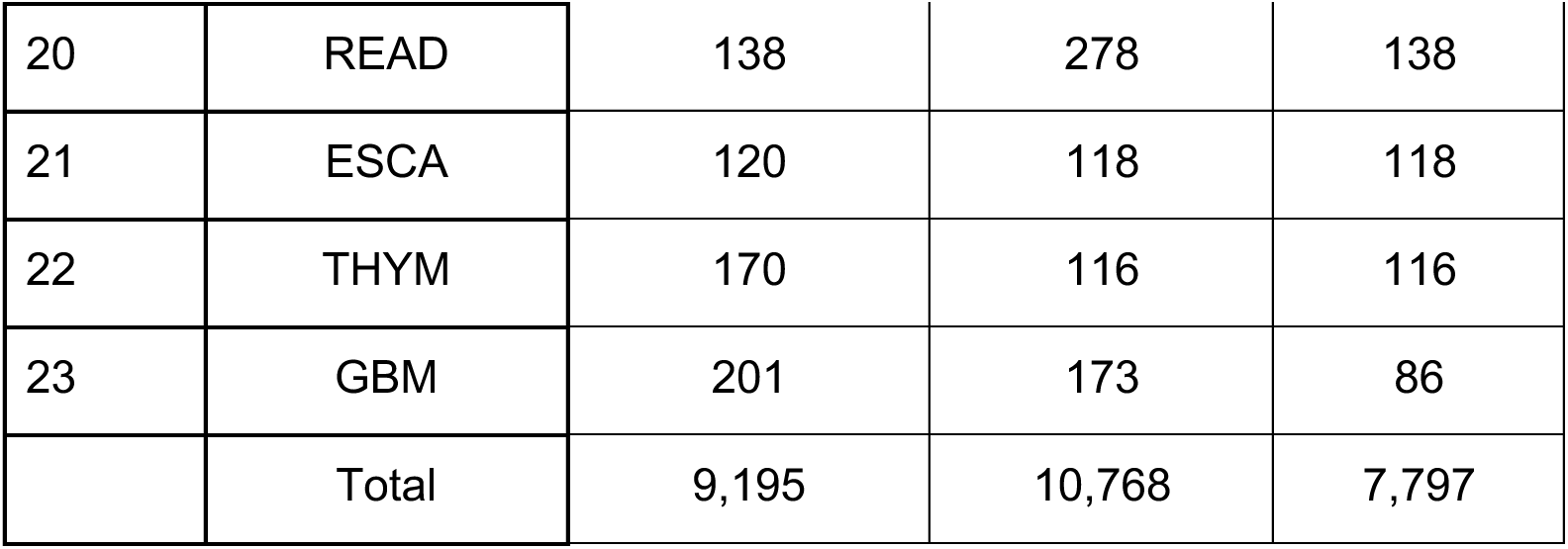
Number of slides and patients used in each TCGA cohort for Path2Omics model development for predicting gene expression.

**Supplementary Table 2.** Number of slides and patients used in each TCGA cohort for Path2Omics model development for predicting DNA methylation.

| No. | Cohort | # FFPE slides | # FF slides | # Patients |
| --- | --- | --- | --- | --- |
| 1 | BRCA | 800 | 951 | 746 |
| 2 | UCEC | 431 | 471 | 387 |
| 3 | THCA | 515 | 529 | 502 |
| 4 | KIRC | 287 | 541 | 287 |
| 5 | LGG | 807 | 644 | 476 |
| 6 | LUSC | 342 | 440 | 329 |
| 7 | LUAD | 436 | 597 | 390 |
| 8 | HNSC | 465 | 645 | 443 |
| 9 | PRAD | 436 | 494 | 390 |
| 10 | BLCA | 447 | 403 | 384 |
| 11 | COAD | 225 | 380 | 221 |
| 12 | LIHC | 377 | 385 | 363 |
| 13 | STAD | 333 | 428 | 308 |
| 14 | KIRP | 263 | 307 | 248 |
| 15 | CESC | 272 | 270 | 262 |
| 16 | SARC | 596 | 254 | 254 |
| 17 | PCPG | 195 | 177 | 177 |
| 18 | PAAD | 202 | 215 | 182 |
| 19 | TGCT | 253 | 155 | 153 |
| 20 | READ | 74 | 138 | 74 |
| 21 | ESCA | 141 | 139 | 139 |
| 22 | THYM | 175 | 121 | 121 |
| 23 | GBM | 87 | 91 | 52 |
|  | Total | 8,159 | 8,775 | 6,888 |

**Supplementary Table 3.** Number of slides and patients in each external cohort for Path2Omics model evaluation for predicting gene expression.

| No. | Cohort | # Slides | # Patients | Slide type |
| --- | --- | --- | --- | --- |
| 1 | NCI-LGG | 309 | 307 | FFPE |
| 2 | CPTAC-BRCA | 106 | 106 | FFPE |
| 3 | CPTAC-KIRC | 312 | 221 | FFPE |
| 4 | CPTAC-LUSC | 109 | 108 | FFPE |
| 5 | CPTAC-LUAD | 224 | 224 | FFPE |
| 6 | CPTAC-COAD | 103 | 103 | FFPE |
| 7 | TransNeo-BRCA | 160 | 160 | FF |
|  | Total | 1,323 | 1,229 |  |

**Supplementary Table 4.** Number of patients in each TCGA cohort for survival analysis based on the predicted gene expression and DNA methylation.

| No. | Cohort | Gene expression | DNA methylation |
| --- | --- | --- | --- |
| 1 | BRCA | 1,022 | 739 |
| 2 | THCA | 495 | 500 |
| 3 | KIRC | 489 | 284 |
| 4 | LUSC | 446 | 320 |
| 5 | LUAD | 426 | 377 |
| 6 | BLCA | 377 | 381 |
| 7 | HNSC | 373 | 393 |
| 8 | COAD | 355 | 211 |
| 9 | LIHC | 333 | 339 |
| 10 | STAD | 263 | 298 |
| 11 | KIRP | 236 | 228 |
| 12 | ESCA | 111 | 132 |
|  | Total | 4,928 | 4,202 |

### Extended Data

**Extended Data Fig. 1.**
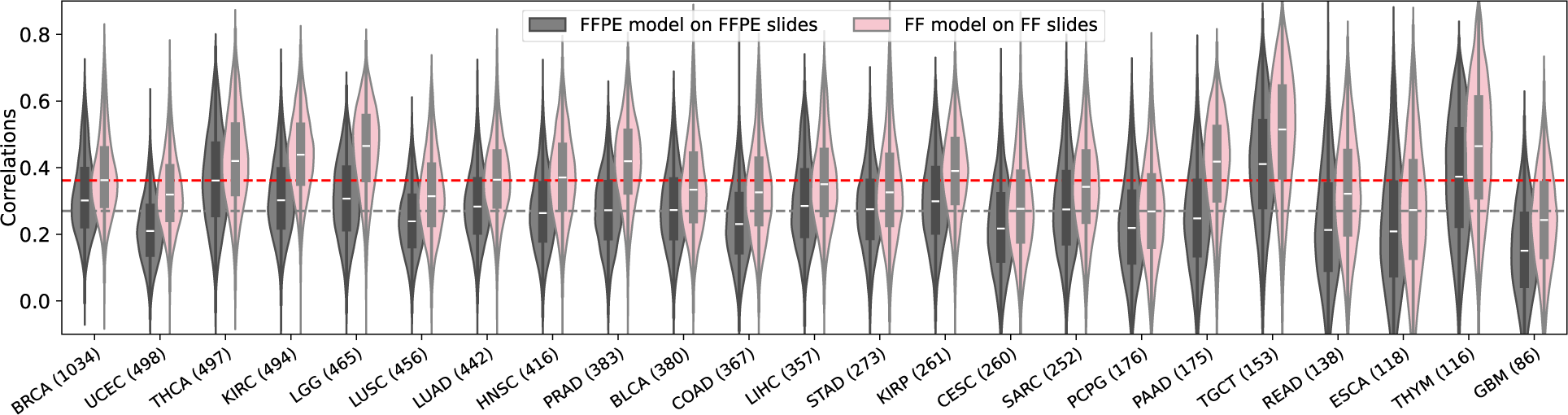
The distribution of correlations between the predicted and actual gene expression values across the cohort samples. The violin plots demonstrate the distribution of Pearson correlations between predicted and measured expression values across the cohort samples for each gene achieved by the Path2Omics-FFPE model on FFPE slides (gray), and the Path2Omics-FF model on FF slides (pink). In the violin plots, the central mark represents the median. The number of patients in each cohort is shown in parentheses.

**Extended Data Fig. 2.**
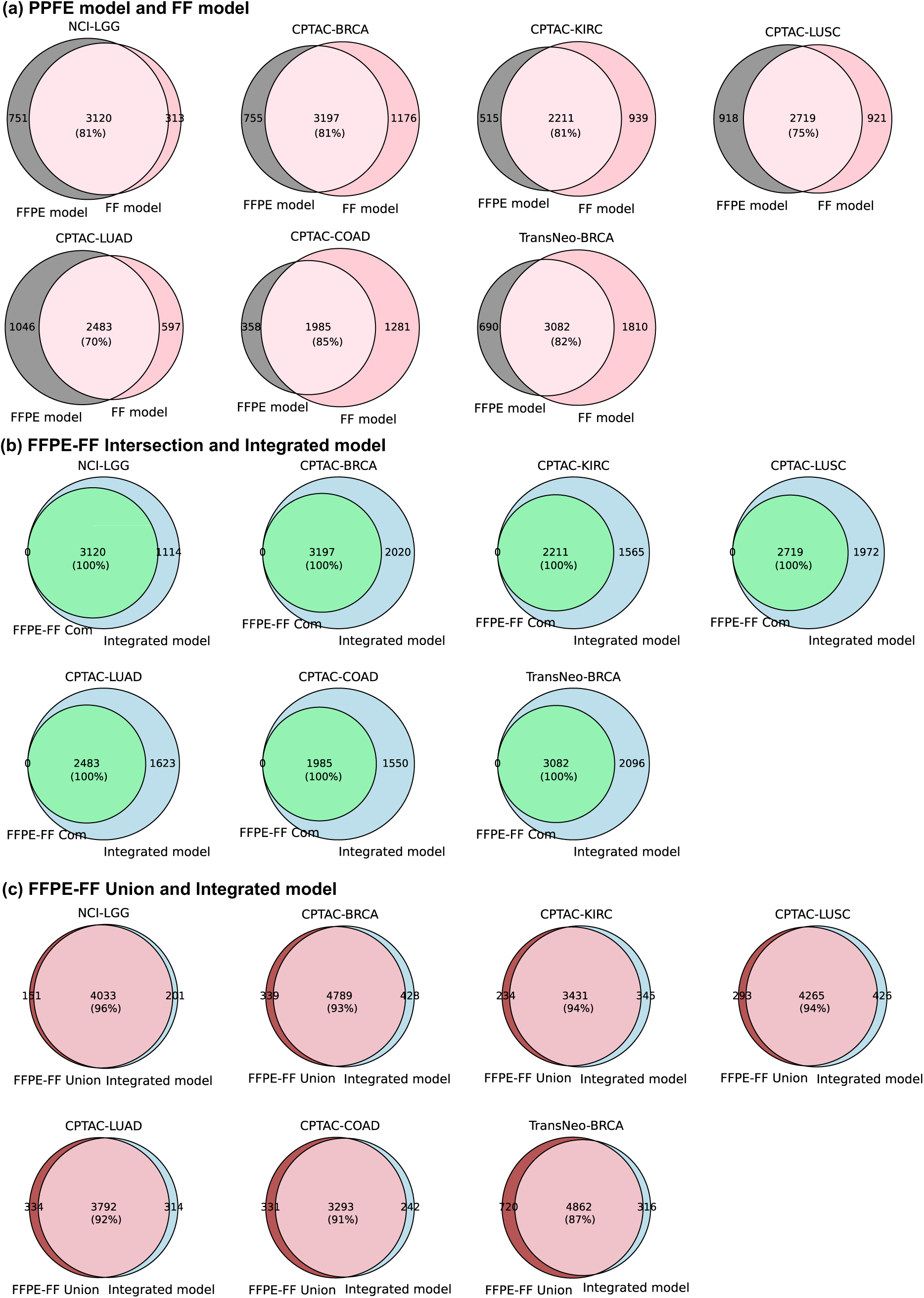
The overlap among the well-predicted genes achieved by each model on external cohorts. (**a**) The Venn diagrams demonstrate the overlap between the well-predicted genes achieved by the FFPE model and the FF model. (**b**) Similar Venn diagrams are shown for the genes that were well-predicted by both the FFPE model and the FF model (FFPE-FF Comm), and the genes that were well-predicted by the Integrated model. (**c**) Similar Venn diagrams are shown for the overlap between the genes that were well-predicted by either the FFPE model or the FF model (FFPE-FF Union) and the genes that were well-predicted by the Integrated model.

**Extended Data Fig. 3.**
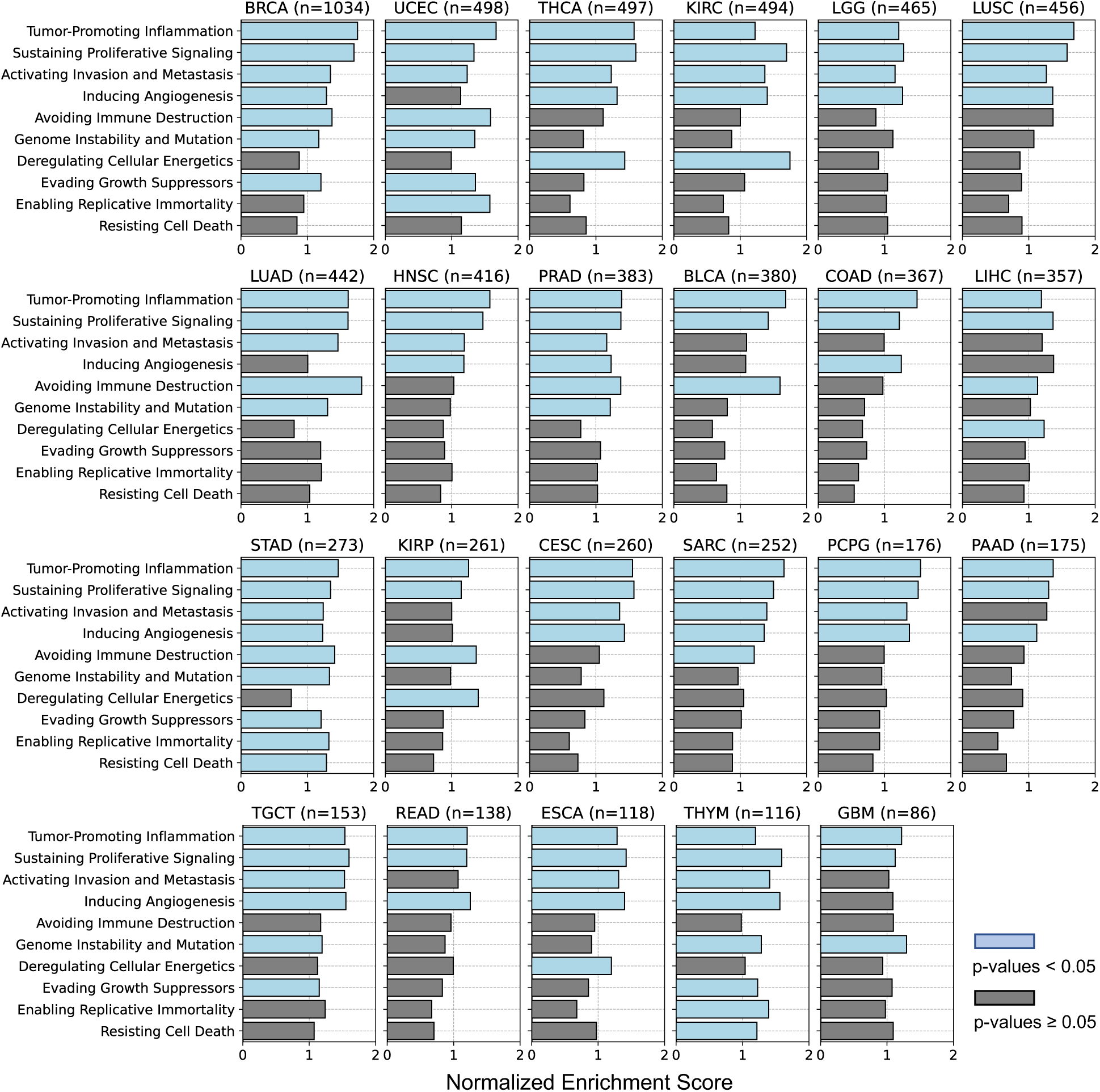
Gene set enrichment analysis identifying pathways associated with the well-predicted genes, achieved by Path2Omics-integrated model on FFPE slides. P-values were calculated using a one-sided permutation test for gene set enrichment analysis. Light blue bars denote significance (p-values < 0.05), gray bars denote non-significance.

**Extended Data Fig. 4.**
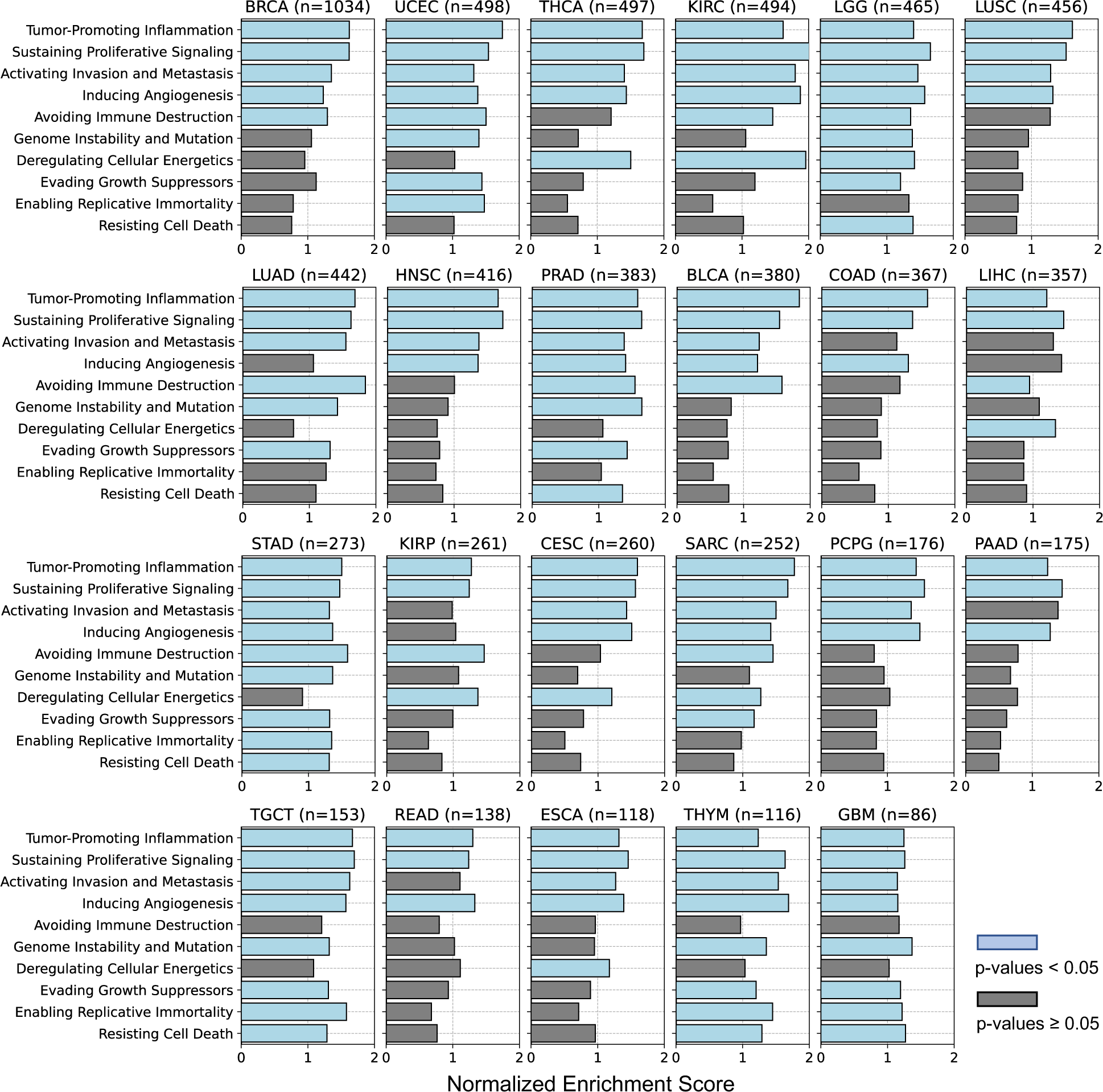
Gene set enrichment analysis identifying pathways associated with the well-predicted genes, achieved by Path2Omics-integrated model on FF slides. P-values were calculated using a one-sided permutation test for gene set enrichment analysis. Light blue bars denote significance (p-values < 0.05), gray bars denote non-significance.

**Extended Data Fig. 5.**
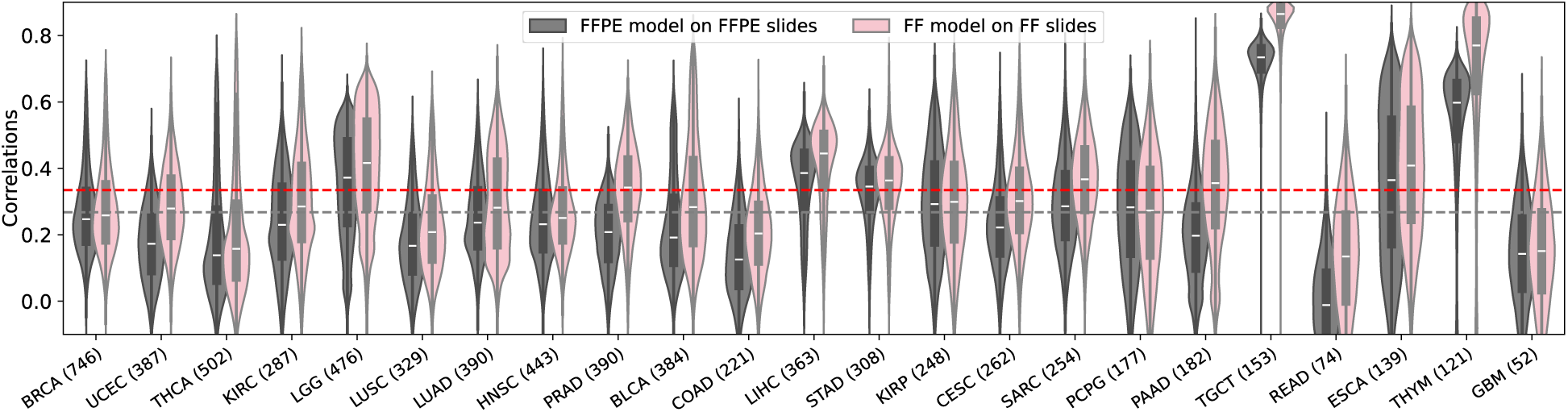
The distribution of correlations between the predicted and actual DNA methylation beta values across the cohort samples. The violin plots demonstrate the distribution of Pearson correlations between predicted and measured DNA methylation beta values across the cohort samples for each CpG site achieved by the Path2Omics-FFPE model on FFPE slides (gray) and the Path2Omics-FF model on FF slides (pink). In the violin plots, the central mark represents the median. The number of patients in each cohort is shown in parentheses.

**Extended Data Fig. 6.**
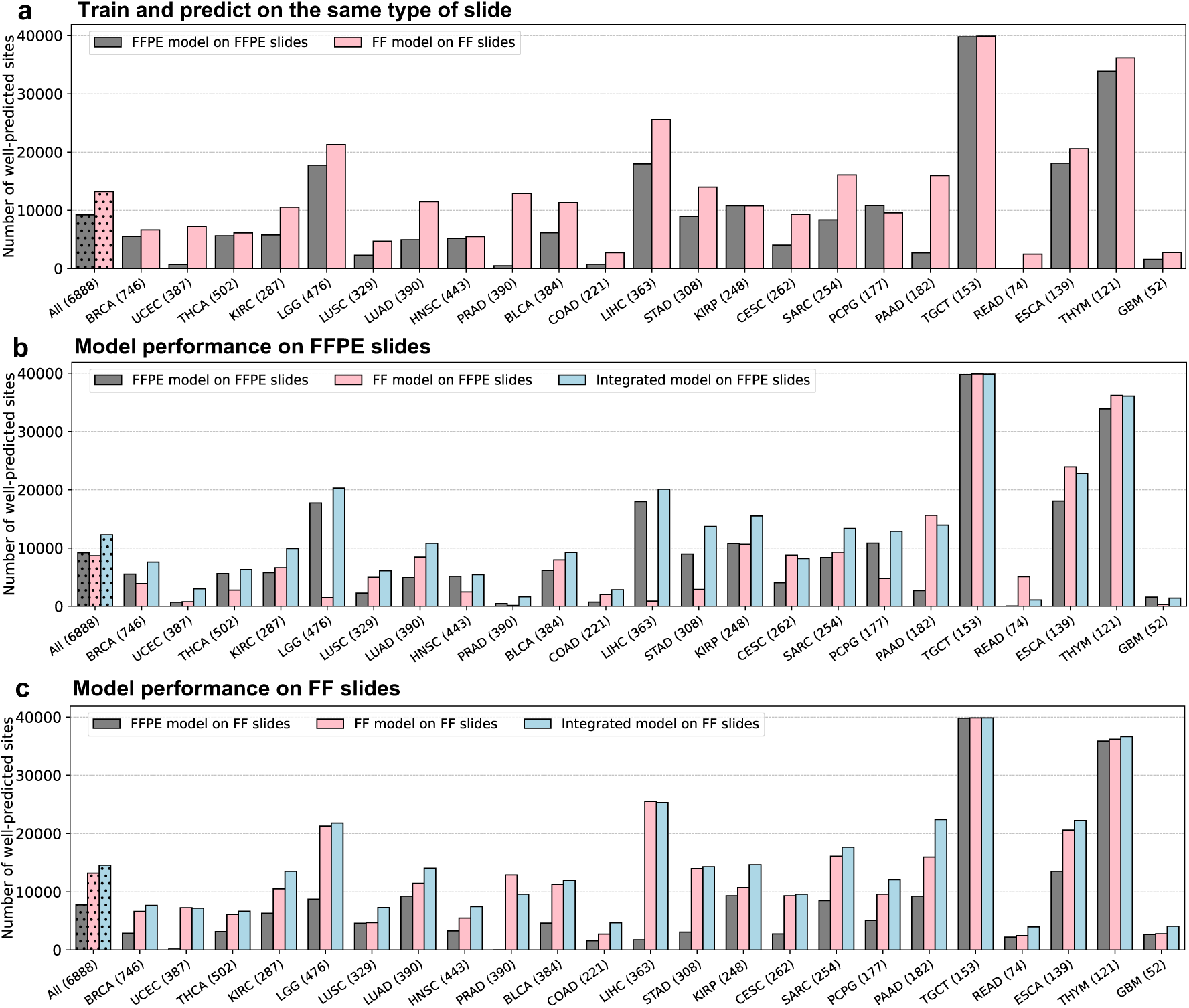
Path2Omics performance in predicting DNA methylation. **(a)** The number of well-predicted CpG sites, defined as having a Pearson correlation between predicted and actual methylation beta values across the cohort samples above 0.4, achieved by the Path2Omics-FFPE model on FFPE slides (gray), and the Path2Omics-FF model on FF slides (pink). The first bars show the average across the 23 cohorts. The number of patients in each cohort is shown in parentheses. (**b)** The number of well-predicted sites archived by the FFPE model (gray), the FF model (pink) and the Integrated model (light blue) when tested on FFPE slides across 23 TCGA cancer cohorts. The last three columns show the average across the 23 cohorts. (**c)** Similar plot to (**b**), but when tested on FF slides.

**Extended Data Fig. 7.**
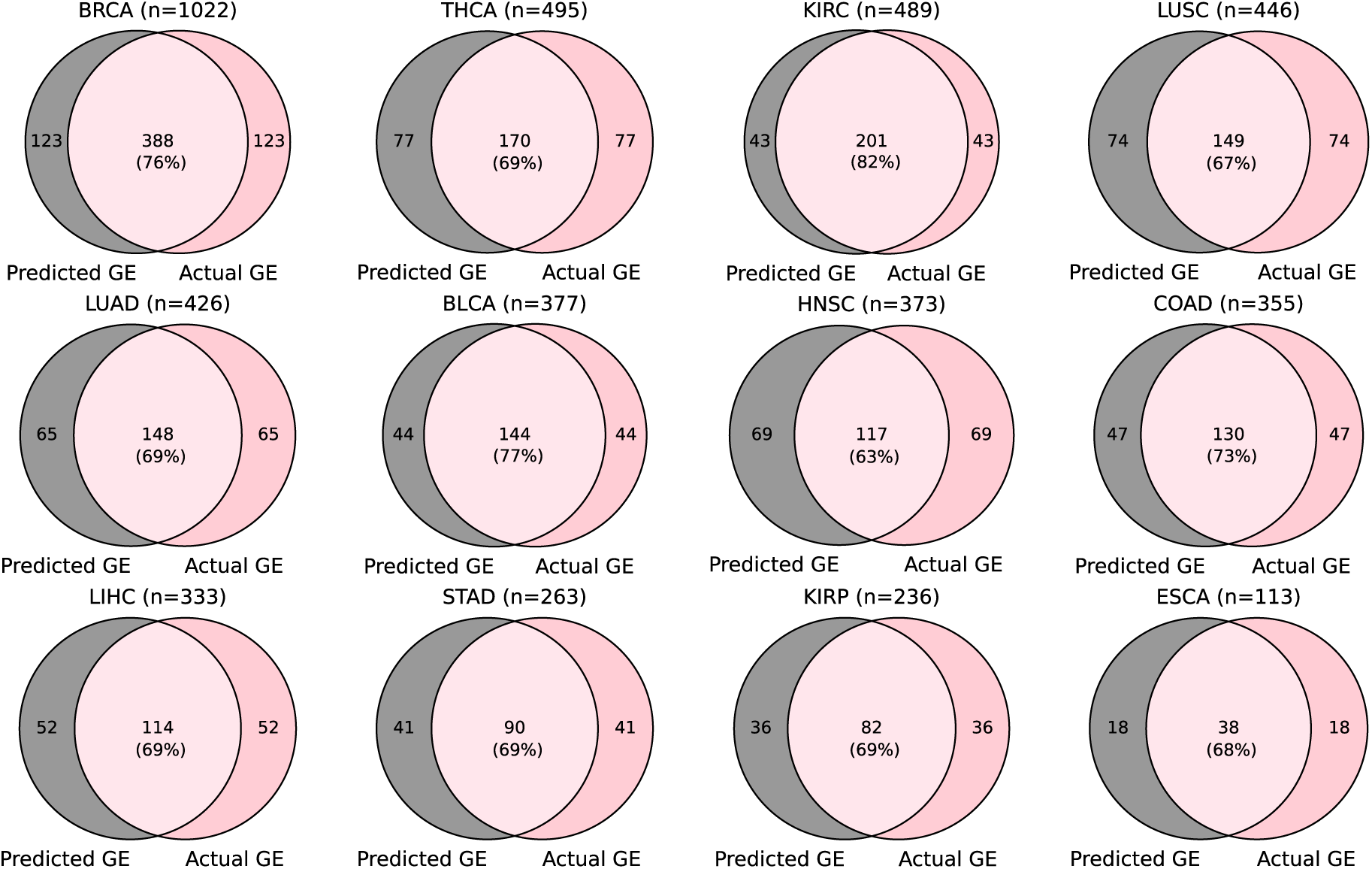
Overlap between patients in high-risk groups assigned by models using predicted (“Predicted GE”, gray) and actual (“Actual GE”, pink) gene expression.

**Extended Data Fig. 8.**
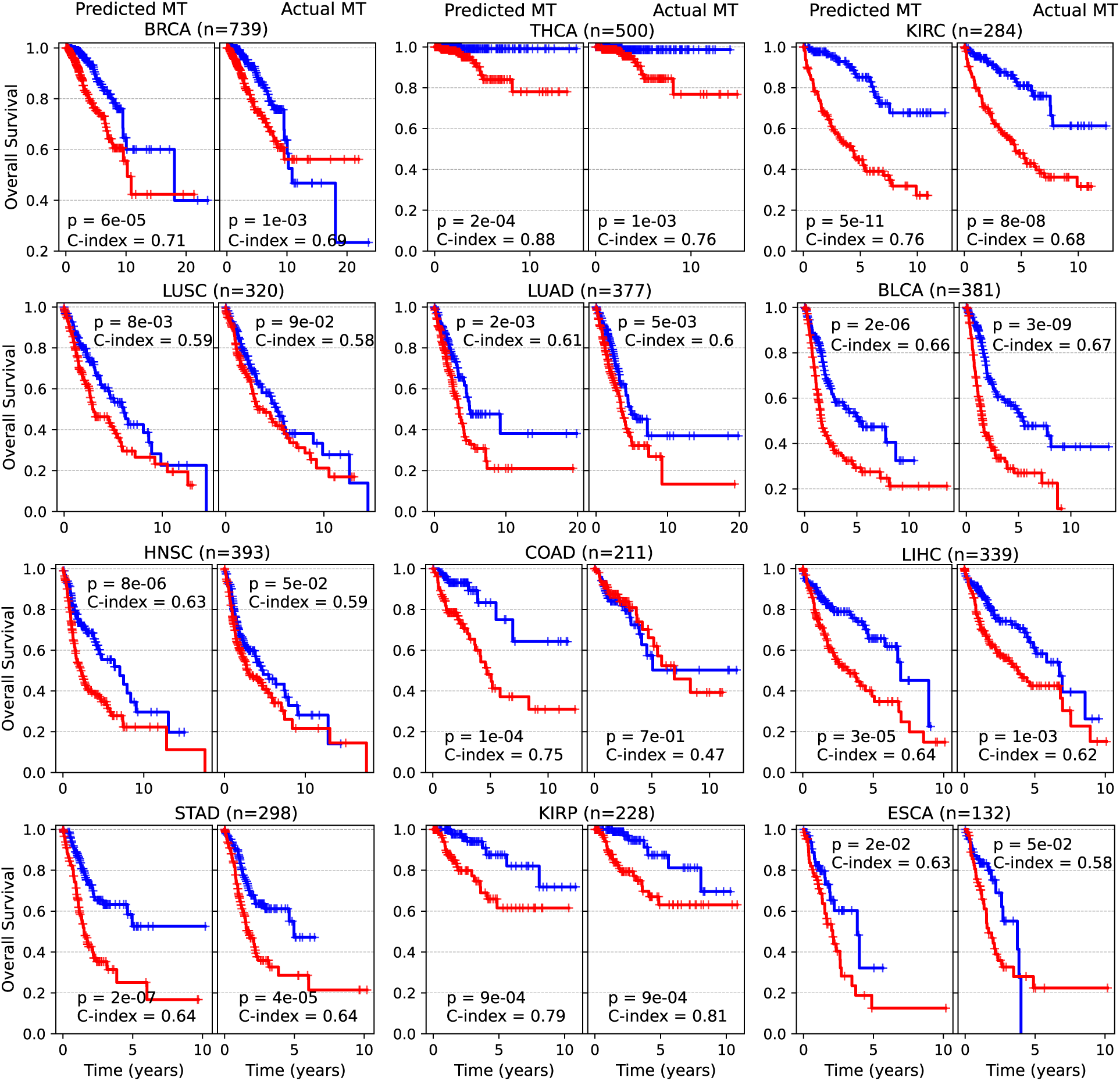
Model performance in predicting patient survival based on the inferred and measured methylation. Kaplan-Meier curves were generated from the model using predicted methylation and patient demographics (sex, age) (“Predicted MT”, left panels) and compared with those from the model using actual methylation and patient demographics (“Actual MT”, right panels) across 12 cancer cohorts. In each cohort, patients were stratified into high-risk (red) and low-risk (blue) groups based on the median risk score generated by each model. P-values were calculated using a two-sided log-rank test.

**Extended Data Fig. 9.**
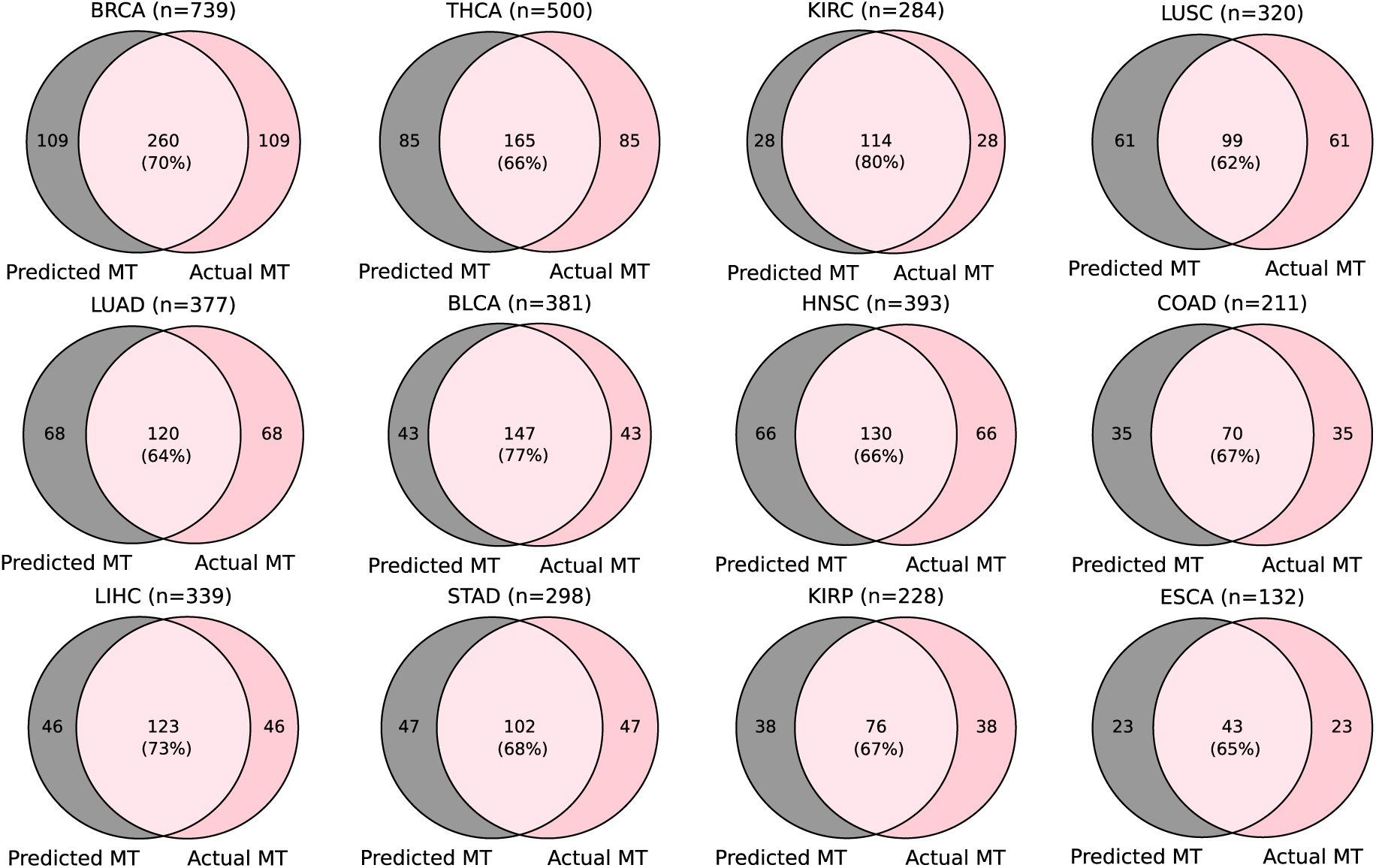
Overlap between patients in high-risk groups identified by models using predicted (“Predicted MT”, gray) and actual (“Actual MT”, pink) methylation.

## References

Alsaafin, Areej, Amir Safarpoor, Milad Sikaroudi, Jason D. Hipp, and H. R. Tizhoosh. 2023. “Learning to Predict RNA Sequence Expressions from Whole Slide Images with Applications for Search and Classification.” Communications Biology 6 (1): 304.

Beck, Andrew H., Ankur R. Sangoi, Samuel Leung, Robert J. Marinelli, Torsten O. Nielsen, Marc J. van de Vijver, Robert B. West, Matt van de Rijn, and Daphne Koller. 2011. “Systematic Analysis of Breast Cancer Morphology Uncovers Stromal Features Associated with Survival.” Science Translational Medicine 3 (108): 108ra113.

Boehm, Kevin M., Emily A. Aherne, Lora Ellenson, Ines Nikolovski, Mohammed Alghamdi, Ignacio Vázquez-García, Dmitriy Zamarin, et al. 2022. “Multimodal Data Integration Using Machine Learning Improves Risk Stratification of High-Grade Serous Ovarian Cancer.” Nature Cancer 3 (6): 723–33.

Bulten, Wouter, Hans Pinckaers, Hester van Boven, Robert Vink, Thomas de Bel, Bram van Ginneken, Jeroen van der Laak, Christina Hulsbergen-van de Kaa, and Geert Litjens. 2020. “Automated Deep-Learning System for Gleason Grading of Prostate Cancer Using Biopsies: A Diagnostic Study.” The Lancet Oncology 21 (2): 233–41.

Chang, P., J. Grinband, B. D. Weinberg, M. Bardis, M. Khy, G. Cadena, M-Y Su, et al. 2018. “Deep-Learning Convolutional Neural Networks Accurately Classify Genetic Mutations in Gliomas.” AJNR. American Journal of Neuroradiology 39 (7): 1201–7.

Cheng, Jun, Jie Zhang, Yatong Han, Xusheng Wang, Xiufen Ye, Yuebo Meng, Anil Parwani, Zhi Han, Qianjin Feng, and Kun Huang. 2017. “Integrative Analysis of Histopathological Images and Genomic Data Predicts Clear Cell Renal Cell Carcinoma Prognosis.” Cancer Research 77 (21): e91–100.

Chen, Mingyu, Bin Zhang, Win Topatana, Jiasheng Cao, Hepan Zhu, Sarun Juengpanich, Qijiang Mao, Hong Yu, and Xiujun Cai. 2020. “Classification and Mutation Prediction Based on Histopathology H&E Images in Liver Cancer Using Deep Learning.” Npj Precision Oncology 4 (1): 1–7.

Chen, Richard J., Tong Ding, Ming Y. Lu, Drew F. K. Williamson, Guillaume Jaume, Andrew H. Song, Bowen Chen, et al. 2024. “Towards a General-Purpose Foundation Model for Computational Pathology.” Nature Medicine 30 (3): 850–62.

Chen, Y., H. Li, A. Janowczyk, P. Toro, G. Corredor, J. Whitney, C. Lu, et al. 2023. “Computational Pathology Improves Risk Stratification of a Multi-Gene Assay for Early Stage ER+ Breast Cancer.” NPJ Breast Cancer 9 (1). 10.1038/s41523-023-00545-y.

Cooper, Lee Ad, Elizabeth G. Demicco, Joel H. Saltz, Reid T. Powell, Arvind Rao, and Alexander J. Lazar. 2018. “PanCancer Insights from The Cancer Genome Atlas: The Pathologist’s Perspective.” The Journal of Pathology 244 (5): 512–24.

Coudray, Nicolas, Paolo Santiago Ocampo, Theodore Sakellaropoulos, Navneet Narula, Matija Snuderl, David Fenyö, Andre L. Moreira, Narges Razavian, and Aristotelis Tsirigos. 2018. “Classification and Mutation Prediction from Non-Small Cell Lung Cancer Histopathology Images Using Deep Learning.” Nature Medicine 24 (10): 1559–67.

Courtiol, Pierre, Charles Maussion, Matahi Moarii, Elodie Pronier, Samuel Pilcer, Meriem Sefta, Pierre Manceron, et al. 2019. “Deep Learning-Based Classification of Mesothelioma Improves Prediction of Patient Outcome.” Nature Medicine 25 (10): 1519–25.

Falahkheirkhah, Kianoush, Tao Guo, Michael Hwang, Pheroze Tamboli, Christopher G. Wood, Jose A. Karam, Kanishka Sircar, and Rohit Bhargava. 2022. “A Generative Adversarial Approach to Facilitate Archival-Quality Histopathologic Diagnoses from Frozen Tissue Sections.” Laboratory Investigation; a Journal of Technical Methods and Pathology 102 (5): 554–59.

Fu, Yu, Alexander W. Jung, Ramon Viñas Torne, Santiago Gonzalez, Harald Vöhringer, Artem Shmatko, Lucy R. Yates, Mercedes Jimenez-Linan, Luiza Moore, and Moritz Gerstung. 2020. “Pan-Cancer Computational Histopathology Reveals Mutations, Tumor Composition and Prognosis.” Nature Cancer 1 (8): 800–810.

Ghaffari Laleh, Narmin, Hannah Sophie Muti, Chiara Maria Lavinia Loeffler, Amelie Echle, Oliver Lester Saldanha, Faisal Mahmood, Ming Y. Lu, et al. 2022. “Benchmarking Weakly-Supervised Deep Learning Pipelines for Whole Slide Classification in Computational Pathology.” Medical Image Analysis 79 (July):102474.

Hanahan, Douglas, and Robert A. Weinberg. 2011. “Hallmarks of Cancer: The next Generation.” Cell 144 (5): 646–74.

He, Bryan, Ludvig Bergenstråhle, Linnea Stenbeck, Abubakar Abid, Alma Andersson, Åke Borg, Jonas Maaskola, Joakim Lundeberg, and James Zou. 2020. “Integrating Spatial Gene Expression and Breast Tumour Morphology via Deep Learning.” Nature Biomedical Engineering 4 (8): 827–34.

He, Kaiming, Xiangyu Zhang, Shaoqing Ren, and Jian Sun. 2016. “Deep Residual Learning for Image Recognition.” 2016 IEEE Conference on Computer Vision and Pattern Recognition (CVPR). 10.1109/cvpr.2016.90.

Hoang, Danh-Tai, Gal Dinstag, Eldad D. Shulman, Leandro C. Hermida, Doreen S. Ben-Zvi, Efrat Elis, Katherine Caley, et al. 2024. “A Deep-Learning Framework to Predict Cancer Treatment Response from Histopathology Images through Imputed Transcriptomics.” *Nature Cancer*, July. 10.1038/s43018-024-00793-2.

Hoang, Danh-Tai, Eldad D. Shulman, Rust Turakulov, Zied Abdullaev, Omkar Singh, Emma M. Campagnolo, H. Lalchungnunga, et al. 2024. “Prediction of DNA Methylation-Based Tumor Types from Histopathology in Central Nervous System Tumors with Deep Learning.” Nature Medicine 30 (7): 1952–61.

Hu, Jing, Chuanliang Cui, Wenxian Yang, Lihong Huang, Rongshan Yu, Siyang Liu, and Yan Kong. 2021. “Using Deep Learning to Predict Anti-PD-1 Response in Melanoma and Lung Cancer Patients from Histopathology Images.” Translational Oncology 14 (1): 100921.

Iorio, Francesco, Luz Garcia-Alonso, Jonathan S. Brammeld, Iňigo Martincorena, David R. Wille, Ultan McDermott, and Julio Saez-Rodriguez. 2018. “Pathway-Based Dissection of the Genomic Heterogeneity of Cancer Hallmarks’ Acquisition with SLAPenrich.” Scientific Reports 8 (1): 6713.

Johannet, Paul, Nicolas Coudray, Douglas M. Donnelly, George Jour, Irineu Illa-Bochaca, Yuhe Xia, Douglas B. Johnson, et al. 2021. “Using Machine Learning Algorithms to Predict Immunotherapy Response in Patients with Advanced Melanoma.” Clinical Cancer Research: An Official Journal of the American Association for Cancer Research 27 (1): 131–40.

Jose, Laya, Sidong Liu, Carlo Russo, Annemarie Nadort, and Antonio Di Ieva. 2021. “Generative Adversarial Networks in Digital Pathology and Histopathological Image Processing: A Review.” Journal of Pathology Informatics 12 (November):43.

Kanopoulos, N., N. Vasanthavada, and R. L. Baker. 1988. “Design of an Image Edge Detection Filter Using the Sobel Operator.” IEEE Journal of Solid-State Circuits 23 (2): 358–67.

Kather, Jakob Nikolas, Alexander T. Pearson, Niels Halama, Dirk Jäger, Jeremias Krause, Sven H. Loosen, Alexander Marx, et al. 2019. “Deep Learning Can Predict Microsatellite Instability Directly from Histology in Gastrointestinal Cancer.” Nature Medicine 25 (7): 1054–56.

Kim, Randie H., Sofia Nomikou, Nicolas Coudray, George Jour, Zarmeena Dawood, Runyu Hong, Eduardo Esteva, et al. 2019. “A Deep Learning Approach for Rapid Mutational Screening in Melanoma.” bioRxiv. bioRxiv. 10.1101/610311.

Levy-Jurgenson, Alona, Xavier Tekpli, Vessela N. Kristensen, and Zohar Yakhini. 2020. “Spatial Transcriptomics Inferred from Pathology Whole-Slide Images Links Tumor Heterogeneity to Survival in Breast and Lung Cancer.” Scientific Reports 10 (1): 18802.

Liu, Jianfang, Tara Lichtenberg, Katherine A. Hoadley, Laila M. Poisson, Alexander J. Lazar, Andrew D. Cherniack, Albert J. Kovatich, et al. 2018. “An Integrated TCGA Pan-Cancer Clinical Data Resource to Drive High-Quality Survival Outcome Analytics.” Cell 173 (2): 400–416.e11.

Macenko, Marc, Marc Niethammer, J. S. Marron, David Borland, John T. Woosley, Xiaojun Guan, Charles Schmitt, and Nancy E. Thomas. n.d. “A Method for Normalizing Histology Slides for Quantitative Analysis.” Accessed December 15, 2023. https://ieeexplore.ieee.org/document/5193250.

Mobadersany, Pooya, Safoora Yousefi, Mohamed Amgad, David A. Gutman, Jill S. Barnholtz-Sloan, José E. Velázquez Vega, Daniel J. Brat, and Lee A. D. Cooper. 2018. “Predicting Cancer Outcomes from Histology and Genomics Using Convolutional Networks.” Proceedings of the National Academy of Sciences of the United States of America 115 (13): E2970–79.

Monjo, Taku, Masaru Koido, Satoi Nagasawa, Yutaka Suzuki, and Yoichiro Kamatani. 2022. “Efficient Prediction of a Spatial Transcriptomics Profile Better Characterizes Breast Cancer Tissue Sections without Costly Experimentation.” Scientific Reports 12 (1): 4133.

Nasrallah, Maclean P., Junhan Zhao, Cheng Che Tsai, David Meredith, Eliana Marostica, Keith L. Ligon, Jeffrey A. Golden, and Kun-Hsing Yu. 2023. “Machine Learning for Cryosection Pathology Predicts the 2021 WHO Classification of Glioma.” *Med (New York*, N.Y.) 4 (8): 526–40.e4.

Ozyoruk, Kutsev Bengisu, Sermet Can, Berkan Darbaz, Kayhan Başak, Derya Demir, Guliz Irem Gokceler, Gurdeniz Serin, et al. 2022. “A Deep-Learning Model for Transforming the Style of Tissue Images from Cryosectioned to Formalin-Fixed and Paraffin-Embedded.” Nature Biomedical Engineering 6 (12): 1407–19.

Pang, Minxing, Kenong Su, and Mingyao Li. 2021. “Leveraging Information in Spatial Transcriptomics to Predict Super-Resolution Gene Expression from Histology Images in Tumors.” bioRxiv. 10.1101/2021.11.28.470212.

Qu, Hui, Mu Zhou, Zhennan Yan, He Wang, Vinod K. Rustgi, Shaoting Zhang, Olivier Gevaert, and Dimitris N. Metaxas. 2021. “Genetic Mutation and Biological Pathway Prediction Based on Whole Slide Images in Breast Carcinoma Using Deep Learning.” NPJ Precision Oncology 5 (1): 87.

Sammut, Stephen-John, Mireia Crispin-Ortuzar, Suet-Feung Chin, Elena Provenzano, Helen A. Bardwell, Wenxin Ma, Wei Cope, et al. 2022. “Multi-Omic Machine Learning Predictor of Breast Cancer Therapy Response.” Nature 601 (7894): 623–29.

Schaumberg, Andrew J., Mark A. Rubin, and Thomas J. Fuchs. 2018. “H&E-Stained Whole Slide Image Deep Learning Predicts SPOP Mutation State in Prostate Cancer.” bioRxiv. 10.1101/064279.

Schmauch, Benoît, Alberto Romagnoni, Elodie Pronier, Charlie Saillard, Pascale Maillé, Julien Calderaro, Aurélie Kamoun, et al. 2020. “A Deep Learning Model to Predict RNA-Seq Expression of Tumours from Whole Slide Images.” Nature Communications 11 (1): 3877.

Shulman, Eldad D., Emma M. Campagnolo, Roshan Lodha, Thomas Cantore, Tom Hu, Maclean Nasrallah, Danh-Tai Hoang, Kenneth Aldape, and Eytan Ruppin. 2024. “Path2Space: An AI Approach for Cancer Biomarker Discovery Via Histopathology Inferred Spatial Transcriptomics.” bioRxiv. 10.1101/2024.10.16.618609.

Teichmann, Marvin, Andre Aichert, Hanibal Bohnenberger, Philipp Ströbel, and Tobias Heimann. 2022. “End-to-End Learning for Image-Based Detection of Molecular Alterations in Digital Pathology.” Medical Image Computing and Computer Assisted Intervention – MICCAI 2022, 88–98.

Tsou, Peiling, and Chang-Jiun Wu. 2019. “Mapping Driver Mutations to Histopathological Subtypes in Papillary Thyroid Carcinoma: Applying a Deep Convolutional Neural Network.” Journal of Clinical Medicine Research 8 (10). 10.3390/jcm8101675.

Tweel, Jan G. van den, and Clive R. Taylor. 2010. “A Brief History of Pathology: Preface to a Forthcoming Series That Highlights Milestones in the Evolution of Pathology as a Discipline.” Virchows Archiv: An International Journal of Pathology 457 (1): 3–10.

Wang, Chuhan, Adam S. Chan, Xiaohang Fu, Shila Ghazanfar, Jinman Kim, Ellis Patrick, and Jean Y. H. Yang. 2025. “Benchmarking the Translational Potential of Spatial Gene Expression Prediction from Histology.” Nature Communications 16 (1): 1–17.

Wang, Xiangxue, Cristian Barrera, Kaustav Bera, Vidya Sankar Viswanathan, Sepideh Azarianpour-Esfahani, Can Koyuncu, Priya Velu, et al. 2022. “Spatial Interplay Patterns of Cancer Nuclei and Tumor-Infiltrating Lymphocytes (TILs) Predict Clinical Benefit for Immune Checkpoint Inhibitors.” Science Advances 8 (22): eabn3966.

Wang, Xiyue, Sen Yang, Jun Zhang, Minghui Wang, Jing Zhang, Wei Yang, Junzhou Huang, and Xiao Han. 2022. “Transformer-Based Unsupervised Contrastive Learning for Histopathological Image Classification.” Medical Image Analysis 81 (October):102559.

Wang, Xiyue, Junhan Zhao, Eliana Marostica, Wei Yuan, Jietian Jin, Jiayu Zhang, Ruijiang Li, et al. 2024. “A Pathology Foundation Model for Cancer Diagnosis and Prognosis Prediction.” Nature 634 (8035): 970–78.

Wang, Yinxi, Kimmo Kartasalo, Philippe Weitz, Balázs Ács, Masi Valkonen, Christer Larsson, Pekka Ruusuvuori, Johan Hartman, and Mattias Rantalainen. 2021. “Predicting Molecular Phenotypes from Histopathology Images: A Transcriptome-Wide Expression-Morphology Analysis in Breast Cancer.” Cancer Research 81 (19): 5115–26.

Xu, Hanwen, Naoto Usuyama, Jaspreet Bagga, Sheng Zhang, Rajesh Rao, Tristan Naumann, Cliff Wong, et al. 2024. “A Whole-Slide Foundation Model for Digital Pathology from Real-World Data.” Nature 630 (8015): 181–88.

Zeng, Y., Z. Wei, W. Yu, R. Yin, Y. Yuan, B. Li, Z. Tang, Y. Lu, and Y. Yang. 2022. “Spatial Transcriptomics Prediction from Histology Jointly through Transformer and Graph Neural Networks.” Briefings in Bioinformatics 23 (5). 10.1093/bib/bbac297.

Zhang, Daiwei, Amelia Schroeder, Hanying Yan, Haochen Yang, Jian Hu, Michelle Y. Y. Lee, Kyung S. Cho, et al. 2024. “Inferring Super-Resolution Tissue Architecture by Integrating Spatial Transcriptomics with Histology.” Nature Biotechnology 42 (9): 1372–77.

Zhang, Fang, Su Yao, Zhi Li, Changhong Liang, Ke Zhao, Yanqi Huang, Ying Gao, Jinrong Qu, Zhenhui Li, and Zaiyi Liu. 2020. “Predicting Treatment Response to Neoadjuvant Chemoradiotherapy in Local Advanced Rectal Cancer by Biopsy Digital Pathology Image Features.” Clinical and Translational Medicine 10 (2): e110.

Zheng, Yuanning, Marija Pizurica, Francisco Carrillo-Perez, Humaira Noor, Wei Yao, Christian Wohlfart, Kathleen Marchal, Antoaneta Vladimirova, and Olivier Gevaert. 2023. “Digital Profiling of Cancer Transcriptomes from Histology Images with Grouped Vision Attention.” bioRxiv : The Preprint Server for Biology, October. 10.1101/2023.09.28.560068.

